# *C. elegans* septins regulate a subset of sensory neuronal cilia via cell-non autonomous mechanisms in supporting glia

**DOI:** 10.1101/2025.10.15.682306

**Authors:** Emilia Filipczak, Sofia Tsiropoulou, Oliver E. Blacque

**Affiliations:** School of Biomolecular and Biomedical Science, University College Dublin, Belfield, Dublin 4, Ireland

## Abstract

Primary cilia rely on compartmentalisation mechanisms that establish the organelle’s protein and lipid composition. In mammalian cells, septin GTPases are reported to facilitate cilium formation, function and molecular composition by regulating a membrane diffusion barrier at the ciliary base transition zone (TZ). Here, we examined septins *in vivo*, within *Caenorhabditis elegans* sense organs. Unexpectedly, loss of one or both septin genes (*unc-59*, *unc-61*) does not cause global defects in cilium structure. Instead, only a subset of ciliated neurons, including the phasmid tail neurons, are affected, with septin mutants displaying short and mispositioned ciliary axonemes due to truncated dendritic processes. Notably, nematode septins do not appear to function at the TZs, with mutants retaining normal gating function and gating complex (MKS & NPHP modules) localisations. Furthermore, double mutant analyses reveal a lack of septin gene interaction with the gating pathways. Strikingly, cell-specific rescue experiments show that UNC-61 regulates phasmid neuronal cilia via a cell non-autonomous mechanism, within supporting glial cells. In addition, UNC-61 localises close to sensory pore adherens cell junctions, and we provide evidence that septin loss disrupts their integrity. Together, our data uncovers an unexpected cell non-autonomous function for glial septins in regulating a subset of sensory neuronal cilia, via a mechanism that may involve cell-cell junctions.

## INTRODUCTION

Primary cilia are microtubule-based structures on the surface of most eukaryotic cell types that sense many environmental stimuli (eg. light, odorants, taste compounds) via signalling receptors and transducers enriched within their membrane and cytosol (Satir, Pedersen and Christensen, 2010). Primary cilia are also essential for tissue homeostasis and development, facilitating cell-cell communication signaling pathways such as those involving Sonic Hedgehog and Wingless (Mill, Christensen and Pedersen, 2023). Templated from a mother centriole-derived basal body, the canonical primary cilium adopts a cylindrical arrangement of nine doublet microtubules enveloped by an extension of the plasma membrane. Despite sharing membrane and cytosol with the rest of the cell, cilia are highly compartmentalised, possessing a unique molecular composition and signaling environment. Such compartmentalisation is achieved in part via diffusion barriers at their base, as well as dedicated molecular trafficking (eg. intraflagellar transport, LIFT) and ectosomal pathways that deliver and remove ciliary proteins (Taschner and Lorentzen, 2016; Jensen and Leroux, 2017; Luxmi and King, 2022; Moran *et al*., 2024). Defects in cilia lead to 20+ clinically distinct and overlapping inherited diseases, or ciliopathies, that collectively affect many tissues and organs (Waters and Beales, 2011). Common pathologies include retinal degeneration, cystic kidneys, bone abnormalities, nervous system malformations, and obesity, many of which co-occur in ciliopathy syndromes such as Joubert Syndrome, Bardet-Biedl syndrome (BBS) and Meckel syndrome (MKS). More than 200 ciliopathy genes are known, many of which function within ciliary compartmentalising pathways (Reiter and Leroux, 2017).

The transition zone (TZ) subcompartment at the ciliary base serves key roles in the organelle’s compartmentalisation. Typically comprising the proximal-most 0.2-1.0 μm of the ciliary axoneme, the TZ is thought to house membrane and size-dependent cytosolic diffusion barriers that limit or ‘gate’ free exchange of proteins and possibly lipids with the rest of the cell (Park and Leroux, 2022; Moran *et al*., 2024). More than 20 ciliopathy proteins are TZ-localised, most within biochemically and/or genetically defined assemblies such as the NPHP (Nephronophthisis), CEP290 and MKS modules. TZ ciliopathy protein loss disrupts cilium formation and structure, including the membrane-microtubule Y-linkers and the ciliary necklace membrane specialisation that defines the TZ; loss of these proteins also disrupts ciliary gating, allowing non ciliary proteins to accumulate within the organelle (Reiter and Leroux, 2017; Park and Leroux, 2022; Moran *et al*., 2024; Moye *et al*., 2025). Additional contributors to ciliary gating at the TZ or within the immediately proximal basal body (BB) region include various nucleoporins and BB distal appendage proteins such as TALPID3, ANKRD26, FBF1 and HYLS-1 (Takao and Verhey, 2016; Wei *et al*., 2016; Yan *et al*., 2020).

In 2010, a mammalian septin protein (SEPT2) was reported to localise at the ciliary base of cultured cells, forming a membrane diffusion barrier required for ciliary signaling (Hu *et al*., 2010). Additional cell culture and zebrafish studies have since shown that several septins reside within the ciliary axoneme and base, where they regulate ciliary length, signaling and the TZ localisations of various gating module proteins (eg. B9D1, TMEM67, TMEM231, CEP290, RPGRIP1L) (Chih *et al*., 2011; Ghossoub *et al*., 2013; Kanamaru *et al*., 2022; Safavian *et al*., 2023). Septins comprise a family of small GTPases (13 in humans; SEPT1-12 and SEPT14), forming filamentous and non-filamentous oligomers that associate with membranes and the cytoskeleton. Operating within cellular scaffolds and membrane diffusion barriers, septins regulate a diverse range of processes such as cytokin*e*sis, cell migration, and cell signaling (Mostowy and Cossart, 2012; Benoit, Poüs and Baillet, 2023).

Despite clear cilia associations for septins, their mechanisms of action remain poorly understood. Indeed, several unresolved observations question their role within ciliary gating pathways. For example, the extent of septin localisations at the ciliary base is somewhat controversial; also, septin loss causes varied effects on the TZ localisations of gating modules, ranging from mild to severe reductions, increases, and ectopic signals within the ciliary axoneme (Hu *et al*., 2010; Chih *et al*., 2011; Ghossoub *et al*., 2013; Kanamaru *et al*., 2022; Safavian *et al*., 2023). It is also noteworthy that septin-ciliary connections are mostly unexplored outside of cultured cells.

To further interrogate the septin-ciliary relationship we have employed the nematode, *Caenorhabditis elegans*, which is a powerful animal model for investigating conserved cilia biology and ciliopathy mechanisms. *C. elegans* possesses just two septin genes, *unc-59* and *unc-61*, for which a wide range of non-essential functions have been reported, related to germline development, cytokinesis cleavage furrow ingression and abscission, anaphase cortical rotation, lipid droplet biogenesis, mitochondrial fission, and axonal migration (Nguyen *et al*., 2000; Finger, Kopish and White, 2003; Maddox *et al*., 2007; Green *et al*., 2013; Pagliuso *et al*., 2016; Chen *et al*., 2021; Zaatri, Perry and Maddox, 2021; Perry *et al*., 2023). However, cilia-related roles for septins are not reported in *C. elegans*.

In *C. elegans* hermaphrodites, primary cilia are only found on 60 sensory neurons (of 302 total neurons), positioned at the distal end of long dendritic processes in the nematode head and tail. Most nematode cilia are located within environmentally exposed pores formed by supporting glia, whilst others are embedded in the surrounding glia processes or cuticle. *C. elegans* cilia adopt highly varied morphologies ranging from canonical 3-6 μm long rods to much larger branched and wing-shaped structures, and serve a diverse range of chemo-, osmo-, and thermo-sensory functions (Inglis *et al*., 2007).

In this study we show that septin loss affects the structure of only a small subset of sensory cilia (eg. phasmids), with no effect on ciliary transition zone integrity and associated pathways. For phasmid cilia regulation, we find that septins function cell non-autonomously in glia to control neuronal dendrite length and cilium positioning, possibly by establishing or maintaining glia-associated cell junctions.

## MATERIALS AND METHODS

### *C. elegans* strains and maintenance

All strains were cultured at 15°C or 20°C on nematode growth medium (NGM) plates seeded with OP50 *E. coli* using standard techniques (Brenner, 1974). To obtain age-synchronised day 1 adults for phenotypic assays, L4 stage hermaphrodite larvae were incubated at 20°C for 16-20 h; alternatively, alkaline hypochlorite treatment of gravid hermaphrodites was used to isolate embryos, which were subsequently incubated at 20°C for ∼65–70 h. All strains used in this study are in **Tables S1 and S2**.

### Generation of *dlg-1::mScarlet* and *unc-61::gfp* knock-in strains

CRISPR Cas 9 protocols were carried out as described in (Paix *et al*., 2015), using a *dpy-10* co-CRISPR approach. The repair templates for GFP and mScarlet (codon optimised for *C. elegans*), including ∼35 bp homology arms, were generated via two rounds of high-fidelity PCR. The double-stranded repair templates were then digested into single stranded donor strands as described (Eroglu, Yu and Derry, 2023) to increase efficiency. All CRISPR protocols used Alt-R Cas9 nuclease 3NLS (Integrated DNA Technologies; IDT, #1074181), Alt-R tracrRNA (IDT, #1072533) and custom-generated gene-specific Alt-R crRNA (IDT). Gene-specific crRNA sequences were selected based on proximity to the desired edit. RNA was reconstituted with 5 mM Tris-HCl pH 7.5 and kept at −80°C. Each injection mix contained 1 μl gene-specific crRNA (0.3 nmol/μl), 1 μl tracrRNA (0.425 nmol/μl), 0.2 μl *dpy-10* crRNA (0.6 nmol/μl), 0.23 μl *dpy-10* ssODN (500 ng/μl), 5-6 μl GFP or wrmScarlet repair template (up to a maximum final concentration of 500 ng/μl), 1-2 μl 1 M KCl, 0.38 μl HEPES (200 mM, pH 7.4), 0.2 μl Cas9 (61 μM), for a final volume of 10 μl. The mixes were prepared on ice, mixed gently, centrifuged at ∼15,000x g for 2 mins and incubated at 37°C for 15 mins prior to injection. F1 progeny was then screened for the desired edit using PCR (primers in **Table S3**). Sanger sequencing was used to verify edit accuracy.

### Generation of strains expressing *unc-61::gfp* as a transgene in phasmid neurons and glia

*unc-61* transgene constructs were generated by fusion PCR (Hobert, 2002). Briefly, *unc-61::gfp* genomic DNA was amplified from a CRISPR-generated knock-in strain (OEB1107) and fused to 5’ UTR sequence of *mir-228* (2.2kb) or *srb-6* (1.5kb) to drive specific expression in glia and phasmid neurons, respectively (Troemel *et al*., 1995; Pierce *et al*., 2008). The linear fusion constructs were injected into WT hermaphrodites at ∼ 1 ng/μL, along with 100 ng/μL of pRF4 [*rol-6(su1006)*] as a co-injection marker, to generate transgenic worms harbouring extrachromosomal arrays. All primers are in **Table S3.**

### Dye-filling assay

Dye-filling assays were performed as previously described in (Sanders, Kennedy and Blacque, 2015). Briefly, worms were incubated in a 1:100 dilution of DiO or DiI (Invitrogen) in M9 for 30 min, and subsequently recovered for 30 min on seeded NGM plates. Worms were then mounted on 4% agarose pads with 40 mM tetramisole and imaged with a wide-field Leica DM5000B epifluorescence microscope. Assays were conducted in triplicate.

### Wide-field epifluorescence imaging and analysis

Young adult worms were anaesthetised in 40 mM tetramisole and mounted onto 4% or 10% agarose pads. Images were taken using a Leica DM5000B upright epifluorescent microscope with a Andor iXon+ camera run by Andor IQ software and collected using x100, x63 (oil immersion) and x40 objectives. All images were analysed using FIJI/ImageJ (NIH). Dendrite and cilia lengths were measured from maximum projections using the segmented line tool. Fluorescence levels of GFP-tagged proteins at the phasmid neuron ciliary transition zones were measured from a single focal plane where both TZs were in focus. A 40×40 pixel box was drawn around the TZs and integrated signal density was measured; the box was then increased by one pixel in each direction and the measurement repeated to calculate (and subtract) the background signal. For colocalisation analysis of UNC-61::GFP and DLG-1::mScarlet, the distance between the GFP signals and mScarlet signals was measured using the segmented line tool. For DLG-1::mScarlet assessment in Figure 4C, each of the two phasmid cilia pairs was classified as either correctly positioned or mispositioned, and then examined for the presence of the 3 areas of DLG-1 signal corresponding to the neuro-glia (sheath), glia-glia (sheath-socket) and glia (socket)-hypodermal cell junctions.

### Airyscan super-resolution imaging

Young adult worms were fixed in ice-cold methanol for ∼ 2 mins, washed with 1x PBS solution and millicule water, and subsequently mounted on poly-lysine coated slides with VECTASHIELD® Antifade mounting medium (Vector Laboratories, Cat. H-1000-10). Slides were then covered with a coverslip and sealed with nail polish. Z-stacks were acquired using a Zeiss LSM800 Airy system with a 63X oil-immersion objective (N.A. = 1.4), and a 488nm laser at 0.2% power. Z-stacks were deconvulved using the ZEN Black software Airyscan processing function, after which maximum projections were derived for data analysis.

### Transmission Electron Microscopy

Young adult hermaphrodites were prepared as previously described (Sanders, Kennedy and Blacque, 2015). Briefly, worms were fixed in 2.5% glutaraldehyde in Sørensen’s phosphate buffer (0.1 M, pH 7.4) for 48 hours at 4°C, followed by post-fixation in 1% osmium tetroxide for 1 hour. Samples were dehydrated through a graded ethanol series, treated with propylene oxide, and embedded in EPON resin at 60°C for 24 hours. Serial ultra-thin (90 nm) sections of the anterior region were cut using a Leica EM UC6 ultramicrotome, mounted on copper grids, stained with 2% uranyl acetate for 20 minutes and 3% lead citrate for 5 minutes, and imaged using a Tecnai 12 (FEI software) at an acceleration voltage of 120 kV.

### Statistical analysis

All statistical analyses were carried out using GraphPad Prism 8.2.1. A Shapiro-Wilk test was used to determine whether data were normally distributed. One-way ANOVA with Tukey post hoc test was used for normally distributed data, while Mann-Whitney or Kruskal-Wallis test with Dunn’s post hoc were used for non-parametric datasets to determine statistical significance. *P*-values <0.05 were deemed significant.

## RESULTS

### *C. elegans s*eptins regulate cilia and dendrite morphology in a subset of sensory neurons

To assess ciliary roles for *unc-59* and *unc-61*, we employed the *tm1939* and *e228* alleles, respectively. Both alleles incur an early premature stop codon due to a 329-bp frame-shifting deletion (*tm1939*) or a nonsense mutation (*e228*), and are likely null (Nguyen *et al*., 2000) (**Figure 1A**).

**Figure 1.**
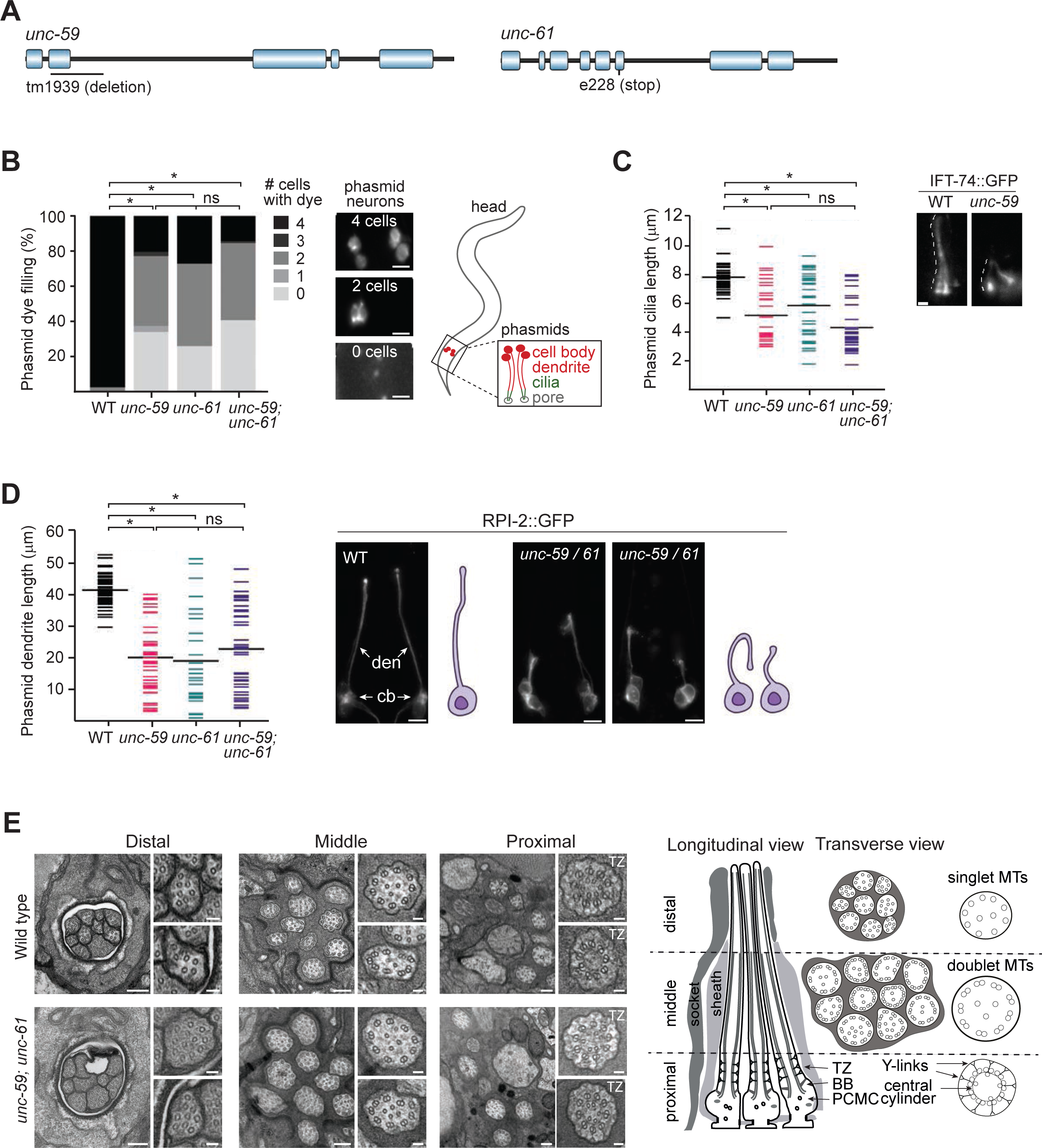
*C. elegans* septins regulate phasmid neuronal cilia and dendrite structure. **(A)** Structure of the two *C. elegans* septin genes, *unc-59* and *unc-61*, showing the location of the deletion (*tm1939*) and nonsense (*e228*) mutations used in this study. **(B)** Phasmid neuron dye-filling (DiI) for single and double septin gene mutants. Histogram quantifies the number of phasmid neurons that fill with dye (0–4). *p<0.0001 (Kruskal-Wallis test with Dunn’s post-hoc analysis vs WT). ns; not significant (p>0.05). Representative fluorescence images of the tail region show animals with four, two, and zero dye-filled phasmid neurons. Scale bars; 10 μm. **(C)** Quantification of phasmid cilia length using an IFT-74::GFP (knock-in) reporter in single and double septin gene mutants. Means indicated by the wide black lines. *p<0.0001 (Kruskal-Wallis test with Dunn’s post-hoc analysis vs WT). ns; not significant (p>0.05). Representative fluorescence images show IFT-74::GFP signals in a phasmid ciliary pair. Scale bar; 1 μm. **(D)** Quantification of phasmid neuron dendrite length using an RPI-2::GFP reporter in single and double septin gene mutants. Means indicated by the wide black lines. Representative fluorescence images show the dendrite length phenotypes in WT and septin gene mutants. Den; dendrite, cell bodies; cb. Scale bars; 10 μm. **(E)** Transmission electron microscopy images showing the ultrastructure of the amphid sensory pore of WT and *unc-59;unc-61* double mutants at the level of the ciliary distal (singlet microtubules), middle and proximal segments. The schematic shows longitudinal and transverse views of a wild type amphid pore (only 3 of 10 axonemes shown in the longitudinal schematic for simplicity). TZ; transition zone, BB; basal body, PCMC; periciliary membrane compartment. Scale bars; 200 nm (large panels); 50 nm (small panels).

First we used a lipophilic dye uptake protocol to assess the integrity of various ciliary subtypes. For this assay, worms are soaked in fluorescent dye (eg. DiO), which incorporates into 6 bilateral amphid (head) and both bilateral phasmid (tail) neurons via their environmentally exposed sensory cilia (Starich *et al*., 1995). Worms with short or absent cilia (eg. IFT gene mutants) are dye-filling defective (Dyf) (Perkins *et al*., 1986; Starich *et al*., 1995). When applied to the septin alleles, we observed a clear Dyf phenotype (loss of uptake in 1 or more neurons) in the majority of phasmid neurons, with no significant difference between single and double mutants (**Figure 1B**). In contrast, amphid neuron dye filling was normal (**Figure S1A**).

As dye filling represents an indirect measure of cilium integrity, cilia were directly visualised using an endogenous (knock-in) IFT-74::GFP reporter, which is expressed in all ciliated sensory neurons (Yi *et al*., 2017). Compared to wild type worms, the phasmid cilia of septin mutants were truncated and often misoriented (**Figure 1C**). Furthermore, these cilia appeared mispositioned, displaced to ectopic anterior regions of the tail, possibly on account of abnormally short dendritic processes. Using an RPI-2::GFP reporter that better illuminates the entirety of the sensory neuron, we confirmed that the phasmid neuronal dendrites of septin mutants are often truncated and misdirected, thereby explaining the cilia mispositioning (**Figure 1D**). In contrast, and in line with the dye-filling observations, the IFT-74::GFP reporter revealed that septin mutants possess a grossly normal amphid channel bundle (10 axonemes), with no evidence of truncated or mispositioned cilia, or malformed dendritic processes (**Figure S1B**). We also examined 2 additional ciliated sensory head neurons employing *str-2*p::GFP (AWC neuron) and *gcy-32*p::GFP (URX neuron) reporters. In each case, dendrite lengths were mostly unaffected in the septin mutants, although the URX dendrite is slightly longer; furthermore; the large AWC wing shaped cilium was also unperturbed (**Figure S1C**).

Finally, we probed the ultrastructure of amphid channel cilia in the septin double mutant using transmission electron microscopy. Like wild type controls, mutant pores contain 10 ciliary axonemes, each displaying normal microtubule number and arrangements, with the characteristic middle (doublet MTs) and distal segment (singlet microtubules) structure intact. The septin mutant also displays normal ciliary transition zone ultrastructure, with clearly defined membrane-microtubule connecting Y-links and an inner apical ring (**Figure 1E**). The only abnormality we detected is that the distal region of one of the mutant amphid pores contains only 9 ciliary axonemes; this defect was observed in the two mutant worms we analysed (**Figure 1E**). As the proximal region of this pore contains the full complement of 10 axonemes, we conclude that the missing axoneme reflects a cilium that is truncated, rather than mispositioned. Thus, one of the 10 amphid channel cilia is abnormally short in septin-disrupted worms.

Together, these data show that septins regulate only a subset of cilia structures in C. elegans.

### *C. elegans* septins do not functionally interact with cilia transition zone gating modules

The cilia subtype-specific defects we observed in septin mutants could be due, at least in part, to abnormalities in ciliary transition zone (TZ) gating, as proposed in several mammalian cell studies (Chih *et al*., 2011; Kanamaru *et al*., 2022; Safavian *et al*., 2023). Indeed, a possible functional connection between *C. elegans* septins and the MKS and NPHP modules is also suggested by the malformed phasmid neuron dendrite phenotype observed in mutant alleles (Williams *et al*., 2008, 2011; Williams, Masyukova and Yoder, 2010).

To investigate if *C. elegans* septins function within TZ pathways, we first investigated if septins are required for the TZ localisation of ciliary gating proteins. Specifically, we examined one MKS (JBTS-14) and one NPHP (NPHP-4) module protein, using GFP-tagged knock-in reporter alleles (Lange *et al*., 2021). Qualitative analysis of phasmid neurons in *unc-59* single and *unc-59;unc-61* double mutants revealed grossly normal TZ localisations and distributions for mNG::JBTS-14 and mNG::NPHP-4, despite the aforementioned cilia misplacement phenotype (**Figure 2A**). In the case of NPHP-4, the transition fiber localisation that lies immediately proximal to the TZ region of wild type worms was also evident in the septin mutants (**Figure 2A**). JBTS-14 and NPHP-4 localisations also appeared normal in the amphid neurons of mutant animals (**Figure 2A**).

**Figure 2.**
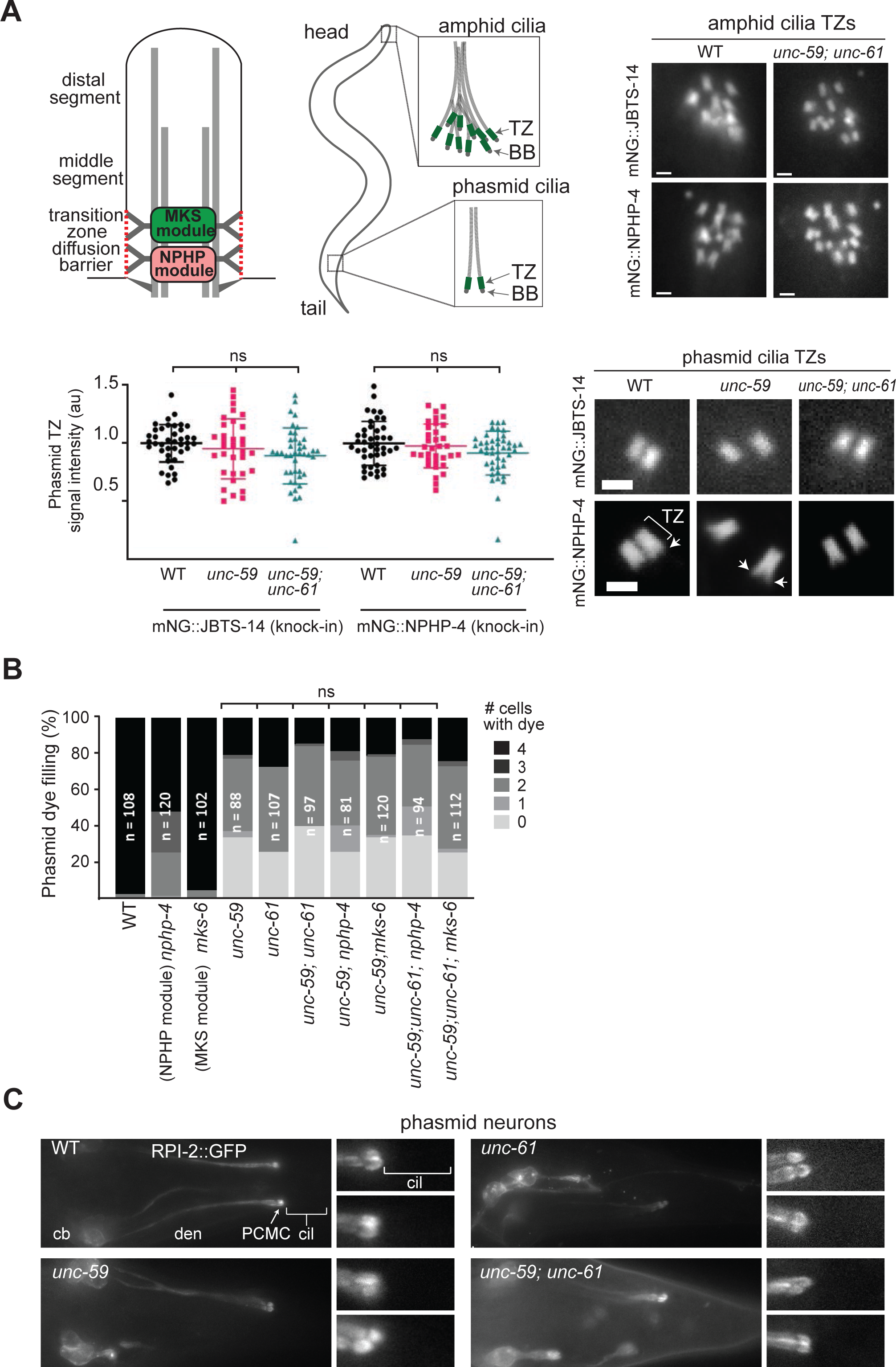
*C. elegans* septins do not appear to functionally interact with TZ gating modules. **(A)** Assessment of mNG::JBTS-14 (MKS module) and mNG::NPHP-4 knock-in reporter localisations in amphid and phasmid neurons of single and double septin gene mutants. Graph shows quantification of fluorescent signals at the phasmid neuron ciliary transition zones (TZ), normalised to WT. ns; not significant (p>0.05; vs WT; one-way ANOVA and Tukey post-hoc analysis). Representative fluorescence images show the TZ localisations of the reporters in single and double septin gene mutants. Bracket; transition zone. Arrows; transition fiber region of the basal body (BB). Scale bars; 1 μm. **(B)** Assessment of phasmid neuron dye-filling in single, double, and triple gene mutants (septins, *mks-6* and *nphp-4*). Histogram quantifies the number of phasmid neurons that fill with dye (0–4). n; number of animals assessed. ns; not significant (p>0.05; vs *unc-59*; Kruskal Wallis test with Dunn’s post-hoc analysis). **(C)** Assessment of the RPI-2::GFP diffusion barrier at the base of cilia. Shown are representative fluorescence images of RPI-2::GFP localisations in the phasmid neurons of single and double septin gene mutants. cb; cell bodies, den; dendrite, PCMC; periciliary membrane compartment, cil; cilia.

Next, we assessed if septin genes genetically interact with the MKS and NPHP modules, which function redundantly with each other to regulate ciliogenesis (Williams *et al*., 2008, 2011). Specifically, we examined cilium integrity (dye filling) in worms containing septin gene mutation(s) *and* a loss of function mutation in an MKS (CC2D2A/MKS6) or NPHP (NPHP4) module component. Analysis of phasmid dye filling revealed that *mks-6* and *nphp-4* mutations do not enhance the Dyf phenotype associated with *unc-59* and *unc-61* mutations (**Figure 2B**). Thus, septin genes do not functionally interact with MKS and NPHP module genes to regulate dye filling.

Finally, we investigated if *C. elegans* septins are required for the TZ membrane diffusion barrier. For this, we employed the RPI-2::GFP reporter, which is enriched at the periciliary membrane, but excluded from the ciliary membrane, of WT worms (Blacque *et al*., 2005). In MKS or NPHP module disrupted worms, RPI-2::GFP ectopically localises to the ciliary membrane, indicating a defect in ciliary gating (Williams *et al*., 2011). We found that RPI-2::GFP remains excluded from septin mutant cilia, indicating that septin loss does not affect the TZ membrane diffusion barrier for this protein (**Figure 2C**).

Taken together with TEM observations (Figure 1E), these data indicate that TZ structure, composition (presence of MKS/NPHP modules) and gating function (at least for lipidated RPI-2) is retained in septin gene mutants. Our double mutant analysis also indicates that septin genes do not functionally interact with MKS and NPHP module genes.

### UNC-61 functions cell non-automously in glia to regulate phasmid neuronal cilia

Next, we employed transgene complementation assays to investigate whether *C. elegans* septins regulate phasmid cilia via cell-autonomous or cell non-autonomous mechanisms. We hypothesised that septins may function in the phasmid neurons or in the supporting phasmid glia (sheath, socket) that form the sensory pores. Indeed, single cell gene expression data reports high levels of *unc-61* RNA in the phasmid socket and other glia, with expression in ciliated neurons limited to a handful of cells, which do not include the phasmids (Taylor *et al*., 2021).

Using fusion PCR, we made linear GFP-tagged *unc-61* (wild type) genomic constructs fused to the 5’UTR of *srb-6*, which drives expression in phasmid neurons, or *mir-228*, which drives expression in most or all glia, including the phasmid sheath and socket cells (Troemel *et al*., 1995; Hobert, 2002; Pierce *et al*., 2008; Fung, Wexler and Heiman, 2020). Transgenic worms housing extrachromosomal arrays of these constructs were generated via gonadal microinjection. As expected, the GFP tag confirmed neuronal and glial expression of *unc-61* using the *srb-6* and *mir-228* promoters (**Figure 3**). Interestingly, the neuronally expressed UNC-61 is diffusely distributed in the phasmid neurons, whereas the glia-expressed UNC-61 accumulates at specific subcellular localisations, especially along the glial processes within the phasmid sensory pore region (**Figure 3**). Next, we crossed the transgenes into the *unc-61(e228)* allele and assessed the resultant strains for phasmid cilium integrity using the dye filling assay. When compared to non-transgenic siblings, mutant worms expressing the glia-expressed *unc-61* transgene exhibited strong rescue of the Dyf phenotype, whereas this was not the case for mutants harbouring the neuronally-expressed *unc-61* transgene (**Figure 3**). Thus, UNC-61 appears to act cell non-autonomously, in phasmid glia, to regulate the integrity of phasmid neuronal cilia.

**Figure 3.**
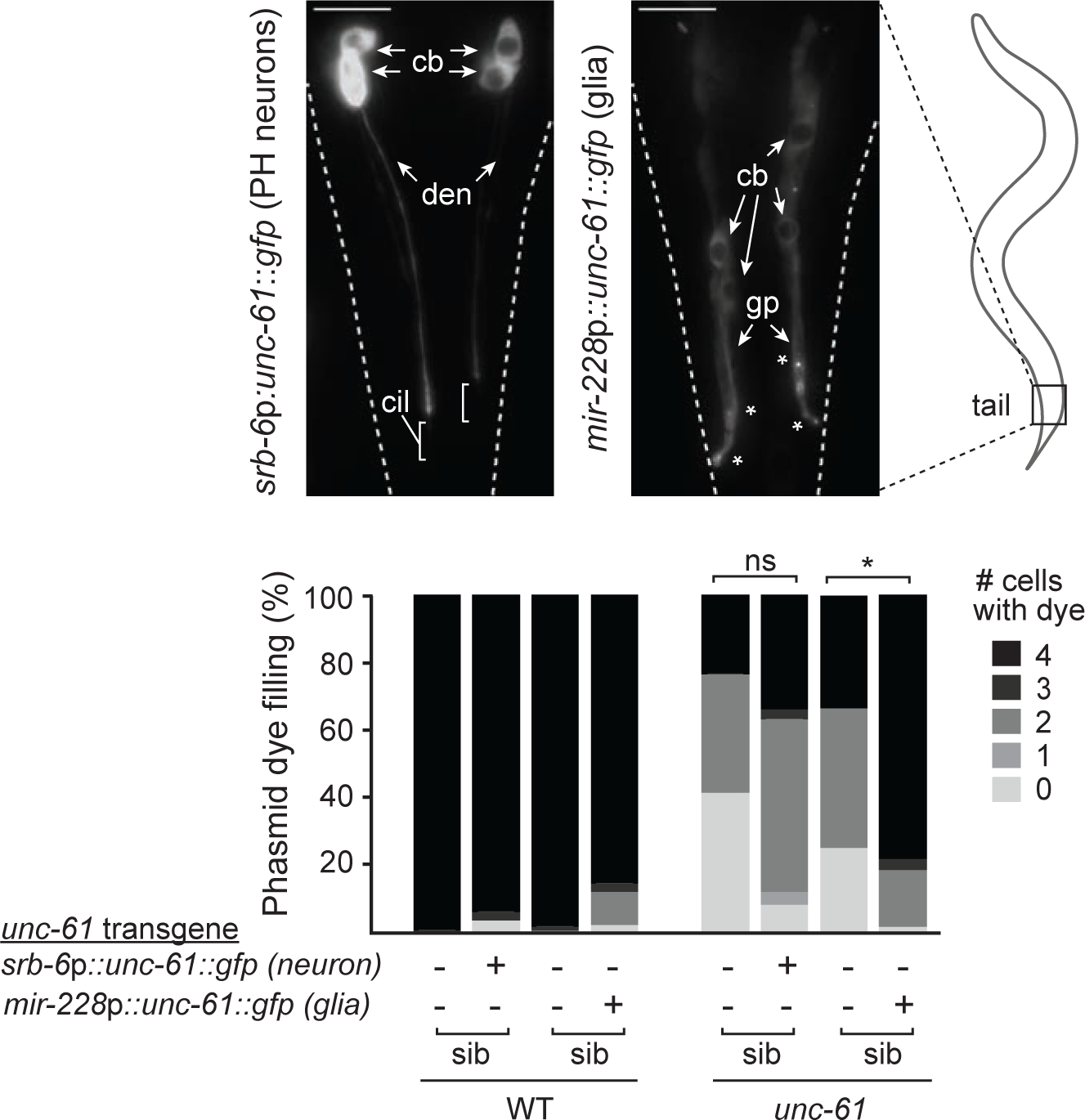
UNC-61 acts cell-non autonomously in glia to regulate cilium integrity. Assessment of phasmid dye filling in WT and *unc-61* mutant worms expressing wild type *unc-61* (GFP-tagged) as a transgene, either in phasmid neurons (via an *srb-6* promoter) or glial cells (via a *mir-228* promoter). Representative fluorescence images of the tail region show UNC-61::GFP localisation in transgenic WT worms. cb; cell bodies; den; dendrite, gp; glial cell process, cil; cilia. Asterisks denote specific localisations near the phasmid pore region. Scale bars; 10 μm. Histogram quantifies the number of phasmid neurons that fill with dye (0–4) in transgenic and non-transgenic siblings (sib), both for WT and *unc-61* mutant worms. *p<0.0001 (Kruskal-Wallis test with Dunn’s post-hoc analysis vs WT). ns; not significant (p>0.05).

### UNC-61 localises at phasmid pore cell junctions

To further investigate the subcellular localisation of *C. elegans* septins in relation to the sensory pores, we used CRISPR-Cas9 gene editing to insert GFP at the 3’ end of the *unc-61* gene. In line with our observations using overexpressed transgenes (Figure 3), endogenously-expressed UNC-61::GFP exhibited several clusters of signal in close proximity to phasmid cilia, marked via TSP-6::mScarlet or DiI (**Figure 4A; Figure S2A**). For day 1 adults, most worms display three discrete clusters, located 1-2 μm proximal to the ciliary axoneme (75% of pores), near the axonemal midpoint (55% of pores), and at the ciliary tip region (100% of pores). The GFP signals typically occur as punctae (1 or 2 per site, very short filaments or rings, with the latter most evident for the distal-most cluster, where UNC-61::GFP clearly wraps around the ciliary tip region (**Figure 4A**).

**Figure 4.**
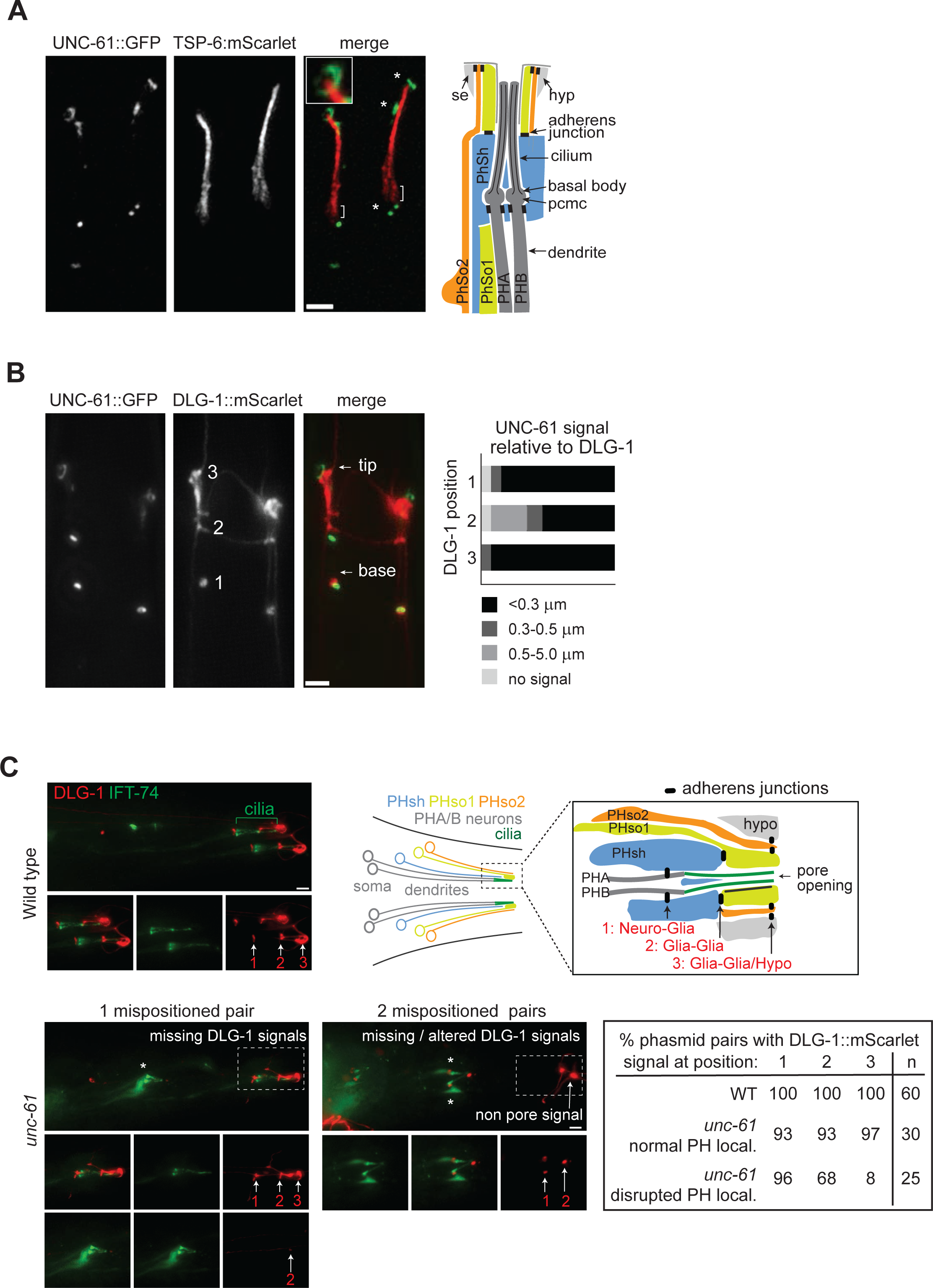
UNC-61 associates with, and is required for, normal adherens cell junctions at the phasmid sensory organ. **(A)** Subcellular localization of endogenously-expressed (knock-in) UNC-61::GFP and TSP-6::mScarlet (ciliary membrane) in the phasmid pore region. Asterisks denote 3 prominent sites of UNC-61::GFP accumulation. Brackets; periciliary membrane compartment. Note that the distal-most UNC-61::GFP signal appears as a ring (inset panel). The schematic shows the organisation of the phasmid sensory pore, comprised of 2 sensory neurons (PHA, PHB), 1 glial sheath cell (PhSh), 2 glial socket cells (PhSo1, PhSo2), and cuticle-associated seam (se) & hypodermal (hyp) cells. Scale bar; 2 μm. **(B)** Subcellular localization of endogenously-expressed UNC-61::GFP and DLG-1::mScarlet (cell junctions) knock-in reporters in the phasmid pore region. Numbers (1, 2, 3) denote 3 prominent DLG-1::mScarlet signals, corresponding to the locations of neuro-glia (1), glia (sheath):glia(socket) and glia (socket):hypodermal/seam cell adherens junctions. Histogram quantifies colocalisation of UNC-61::GFP relative to DLG-1::mScarlet at the 3 prominent positions. UNC-61::GFP signals are binned by distance from DLG-1::mScarlet (<0.3 µm apart; 0.3-0.5 µm apart; 0.5-5.0 µm apart; no UNC-61::GFP signal). Scale bar; 2 µm. **(C)** Assessment of phasmid organ cell junctions in septin disrupted worms. Representative fluorescence images show DLG-1::mScarlet at the phasmid pore region of WT and *unc-61* mutants, with ciliary axonemes visualised using an IFT-74::GFP (knock-in) reporter. Arrows denote the DLG-1::mScarlet adherens cell junction signals (1, 2, 3). The table quantifies the presence or absence of DLG-1::mScarlet signals at positions 1-3. *unc-61* data is split into 2 groups; phasmids that are normally positioned (normal PH local.), and phasmids that are ectopically positioned to anterior regions of the tail (disrupted PH local.). Asterisks denote misplaced cilia. Scale bars; 2 µm.

The position and pattern of these signals is strongly suggestive of an UNC-61 association with cell-cell junctions, which form neuro-glia (sheath), glia-glia (sheath-socket) and glia (socket)-hypodermal contacts at the phasmid sensory pore (see schematic in **Figure 4A**). To explore a possible cell junction localisation for UNC-61, we first established an endogenous mScarlet-tagged reporter for the DLG-1 apical junction protein using CRISPR-Cas9 editing. DLG-1 is a well-characterized scaffolding protein that localizes to adherens junctions and is essential for maintaining epithelial polarity and cell junction integrity in *C. elegans (Firestein and Rongo, 2001)*. As expected, DLG-1::mScarlet is observed throughout the worm, forming punctate and highly elongated signals consistent with the extensive junctions of epithelial cells (McMahon *et al*., 2001) (**Figure 4B; Figure S2B**). In the phasmid sensory pore region, the DLG-1::mScarlet signals are largely punctate and either colocalise with, or are located within 1-2 µm of the three aforementioned UNC-61::GFP signals (**Figure 4B**). In the nosetip region, discrete UNC-61::GFP signal are also very close to DLG-1::mScarlet signals, although the extent and specificity of UNC-61/DLG-1 colocalisation in the context of the anterior sensory pores (eg. amphids) is difficult to determine on account of the widespread nature of the DLG-1 signal (**Figure S2B**).

Together, these findings show that UNC-61 is localised at or very near to cell-cell junctions in the phasmid sensory pore region. However, the extent of colocalisation in the head regions remains inconclusive.

### UNC-61 loss disrupts DLG-1 cell junctions associated with the phasmid sensory organ

As UNC-61 colocalises with DLG-1, we investigated whether septins are required for DLG-1 cell junction integrity by examining DLG-1::mScarlet (knock-in) localisations at the phasmid pores of *unc-61(e228)* mutant worms. The IFT-74::GFP (knock-in) reporter was used to mark the phasmid ciliary axonemes and dendrites. We found that the normally positioned pores in the *unc-61* mutant (∼50 % of all pores) retained all 3 associated DLG-1 signals, namely, neuro-glia (1), glia-glia (2) and glia-hypodermal/seam (3) (**Figure 4C**). For those pores that are abnormal, as denoted by mispositioned ciliary axonemes and short dendrites, the *unc-61* mutant retains apparent neuro-glia DLG-1 signals proximal to the mispositioned cilia (**Figure 4C**). This suggests that the sheath cell and neuronal dendritic processes remain attached and that the sheath cell process may also be truncated. Some DLG-1 signal, reminiscent of a glia(sheath)-glia(socket) junction, is also observed near the tips of mispositioned cilia; thus, the phasmid socket cell 1 (PhSo1) process may also be short in *unc-61* worms (**Figure 4C**). However, for 32% of mispositioned cilia, this presumptive glia-glia DLG-1 signal is not detected near the axonemes. In contrast to the neuro-glia and glia-glia signals, the glia-hypodermal/seam cell DLG-1 signals are dramatically reduced or entirely absent for those *unc-61* mutant phasmids that don’t form properly (ie, where mispositioned cilia are observed) (**Figure 4C**). From these data, we conclude that septin loss affects glia-associated DLG-1 levels at the phasmid sensory organ, especially those associated with hypodermal and seam cells.

## DISCUSSION

Our findings reveal that *C. elegans s*eptins regulate cilium structure, positioning and dendrite extension in a subset of sensory neurons. Specifically, septin gene loss causes severe morphological defects in the phasmid tail neurons, without affecting most examined sensory neurons in the head (amphid channel, AWC, URX), with the exception of one amphid channel cilium. Notably, these defects do not appear to arise from disruption of canonical transition zone (TZ) gating pathways as has been described in mammalian cell culture models (Hu *et al*., 2010; Ghossoub *et al*., 2013; Kanamaru *et al*., 2022). Rather, our cell specific rescue experimentation points to a cell non-autonomous role for septins in glia that support the phasmid sensory neurons. Two observations indicate that this function may relate to cell-cell junctions. First, endogenously tagged reporters reveal that UNC-61 is enriched near or at DLG-1-marked adherens junctions associated with the phasmid pore. Second, septin loss disrupts some DLG-1 localisations within this region. We propose that septins contribute to the structural organisation of glial cell adherens junctions, which in turn, regulate dendritogenesis and ciliogenesis in a subset of sensory neurons.

### Septins regulate only a subset of sensory cilia in *C. elegans*

Thus far, septins have been investigated in cultured mammalian cells and zebrafish, revealing variable and somewhat conflicting roles with regards to cilium formation and length regulation. In early studies, SEPT2, 7 or 9 depletion was reported to reduce ciliation and cilium length in RPE1 and IMCD3 cells (Hu *et al*., 2010; Ghossoub *et al*., 2013). In line with these findings, SEPT9 knockout cells (RPE1) fail to ciliate, and phosphorylation- or methylation-defective variants of SEPT2 and SEPT7 exert dominant negative effects on ciliogenesis in NIH3T3 and RPE1 cells (Toriyama *et al*., 2017; H.-Y. Wang *et al*., 2021; Safavian *et al*., 2023). Furthermore, Sept6 and Sept7 zebrafish morphants show reduced ciliogenesis and short cilia in the Kuppfer’s vesicle and pronephros (Dash *et al*., 2014; Zhai *et al*., 2014). In contrast to these findings, however, is a recent report showing that septin (2/6/7/9)-depleted and knockout (SEPT2) cells (RPE1) possess increased cilia lengths and incidence, suggesting that septins act as negative, rather than positive, ciliogenesis regulators (Kanamaru *et al*., 2022).

Here, using worms where both septin genes (*unc-59*, *unc-61*) are disrupted, we show that septins regulate the formation of a subset of sensory cilia. In mutants, both phasmid tail cilia are frequently short (although sometimes of normal length), and one of the ten ciliary axonemes is missing from the amphid pore channel in the head. However, most analysed cilia appeared morphologically wild-type in septin-disrupted worms. Indeed, electron microscopy reveals that the amphid pore cilia of septin double mutants cilia are grossly normal in terms of microtubule number and organisation within proximal and distal segments. Ciliary membranes, including the large and elaborate AWC ciliary membrane, also appeared normal in mutants. As the mutations employed are strong loss-of-function or null alleles, we conclude that *C. elegans* septins are dispensable for the formation and length regulation of most ciliary subtypes. We were unable to assess the effect of septin gene loss on sensory neuronal function because the *unc-59* and *unc-61* alleles possess a strong uncoordinated (Unc) movement phenotype that prevents the use of chemotaxis, osmolarity sensing and foraging behaviour assays. Thus, it remains to be shown if *C. elegans* septins regulate ciliary functions as has been demonstrated for RhoA and sonic hedgehog signaling in mammalian cells (Hu *et al*., 2010; Kanamaru *et al*., 2022; Safavian *et al*., 2023).

### *C. elegans* septins do not appear to function at the ciliary transition zone

Several mammalian cell culture studies implicate septins as regulators of ciliary gating at the transition zone. Specifically, septin (SEPT2/9) depletion or loss increases mobility of membrane proteins into cilia, and variably disrupts the TZ localisations of gating module proteins (RPGRIP1L, TCTN, MKS3, CEP290, B9D1, CC2D2A, TMEM231) (Hu *et al*., 2010; Chih *et al*., 2011; Kanamaru *et al*., 2022; Safavian *et al*., 2023). Although septins (SEPT2/6/7/9/11) are reported to localise along the ciliary axoneme and/or the ciliary base in mammalian cells and zebrafish, it is debatable if this localisation includes the TZ, which is typically very small (<0.5 micron long) (Hu *et al*., 2010; Ghossoub *et al*., 2013; Dash *et al*., 2014; Fliegauf *et al*., 2014; Zhai *et al*., 2014; Kim *et al*., 2016; Kanamaru *et al*., 2022; Safavian *et al*., 2023). Nonetheless, one study shows SEPT2 localisation at the 1.5 μm long TZ (connecting cilium) of human photoreceptor cells, even though another study reports exclusion of SEPT2 and SEPT7 from the TZ of RPE1 cells (Ghossoub *et al*., 2013; Kanamaru *et al*., 2022). The basis for these differences may relate to cell type specific differences.

Our data indicates that *C. elegans* septin genes do not function at, or regulate, TZ processes. Using endogenous-tagged reporters, we show that septin gene loss does not affect the restricted TZ localisations and levels of MKS (JBTS-14) and NPHP (NPHP-4) module components. Furthermore, epistasis experiments reveal a lack of genetic interaction between septins and gating module genes in regulating cilium integrity (dye-filling); thus, septins do not behave like MKS or NPHP module components. In addition, septin mutants possess intact TZ ultrastructure, and a normal membrane diffusion barrier as assessed via an RPI-2::GFP ciliary exclusion assay. Lastly, using an endogenous GFP-tagged reporter, we show that UNC-61 is excluded from cilia, including the ciliary transition zone and base. These results argue against a direct interaction between septins and the TZ complex in worms. Instead, we propose an alternative mechanism for the ciliogenesis defects observed in septin mutants, involving glia-derived support of sensory neuron morphogenesis, as discussed below.

### *C. elegans* septins control sensory organ morphogenesis

Our finding of highly truncated phasmid dendrites and mispositioned/malformed cilia in *unc-59* and *unc-61* mutants implicates septins as regulators of phasmid sense organ morphogenesis. In *C. elegans*, most ciliated sensory organs contain glial cell (sheath, socket) processes that wrap around the ciliary axonemes to form environmentally exposed pores. For example, in the amphid sense organ, a sheath cell process wraps round the proximal portion of the ciliary axonemes, connecting to each neuron via adherens junctions at the base of the perciliary membrane compartment; a second glial process, from the socket cell, envelops the sheath and the distal ciliary regions, forming adherens junction contacts with the sheath and the adjoining hypodermal cell (Ward *et al*., 1975; Perkins *et al*., 1986; Doroquez *et al*., 2014; Low *et al*., 2019) (see schematics in **Figures 4, 5**). Not all sensory cilia are ensheathed by two glia; for example, several amphid cilia terminate solely within the amphid sheath, and the URX and BAG neuronal endings lack glia ensheathment entirely, although they form anchoring contacts with the IL2 socket cell (Ward *et al*., 1975; Perkins *et al*., 1986; Doroquez *et al*., 2014; Low *et al*., 2019; Cebul, McLachlan and Heiman, 2020; Heiman and Bülow, 2024). Although not yet well understood, sense organ formation involves initial ensheathment of nascent dendritic endings by emerging glial cell processes; subsequently, the cell bodies migrate, resulting in trailing deposition of mature neuronal and glial processes via a retrograde extension mechanism (**Figure 5**) (Heiman and Shaham, 2009; Low *et al*., 2019; Cebul, McLachlan and Heiman, 2020). In terms of timing, nascent dendritic endings are first wrapped by the sheath process, after which the socket process completes glial tube formation (Wexler, Kolotuev and Heiman, 2025). At the onset of cell migration, cilia have yet to form, at least in the amphid sense organ (Low *et al*., 2019).

**Figure 5.**
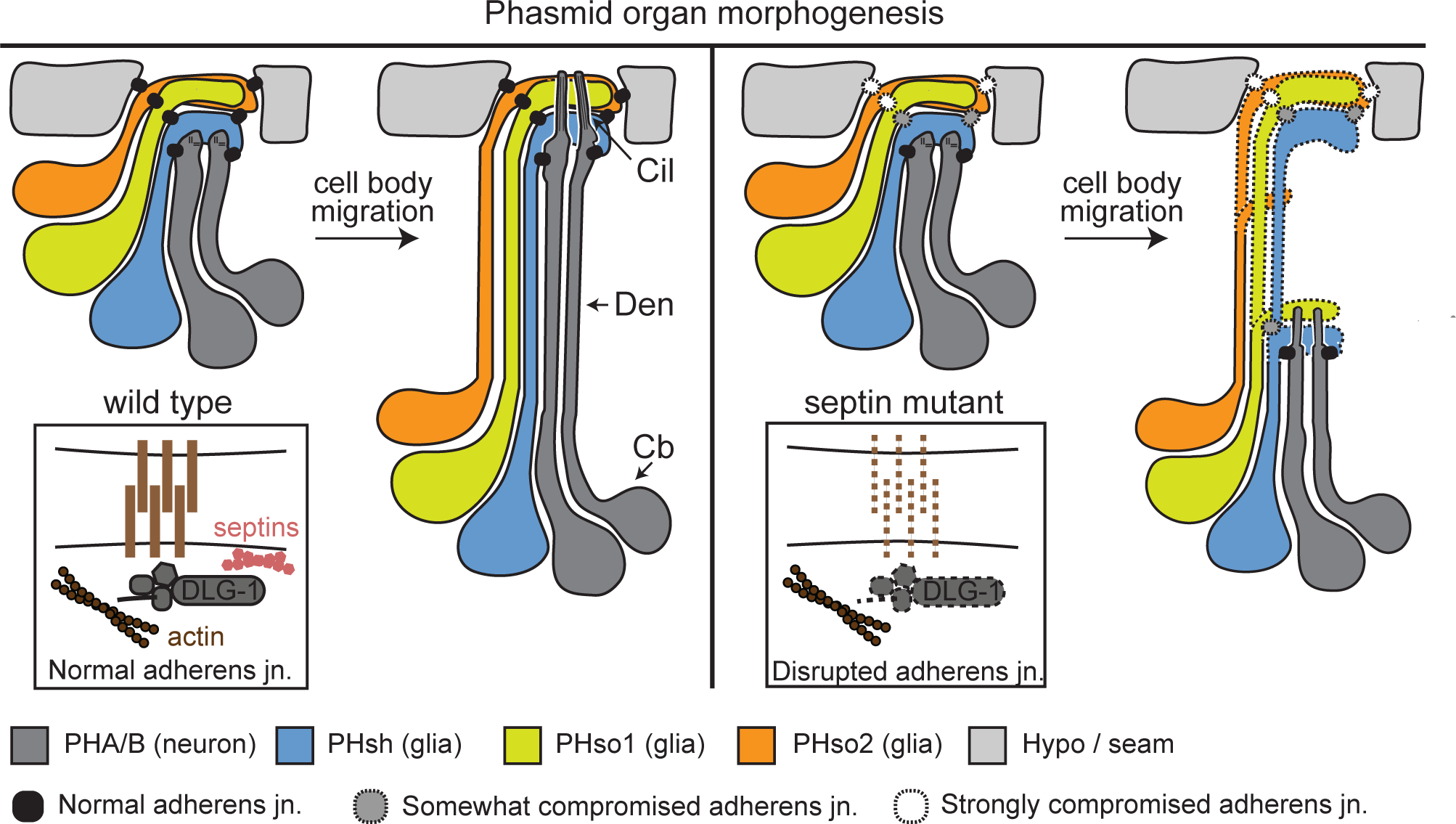
Model schematic. During WT phasmid sensory pore development, adherens cell junctions (neuro-glia, glia-glia, glia-hypodermal/seam) anchor nascent cell processes during neuronal and glial cell body migration. Anchoring facilitates dendrite elongation via a retrograde extension model (Heiman and Shaham, 2009), ensuring correct placement of the sensory pore, and normally elongated ciliary axonemes. Septins, expressed in glia, are proposed to contribute to the stability of DLG-1-positive cell junctions. In this model, loss of junctional integrity weakens cell process anchoring, resulting in truncated neuronal dendrites (and possibly glial processes; denoted by dotted lines), as well as short/misplaced cilia, as the cell bodies migrate. Septin loss may differentially affect the cell junctions, with the most severe effects on the glia-hypodermal/seam cell junctions.

Several regulators of retrograde dendritic extension have been identified. These include the secreted tectorin-like proteins, DEX-1 and DYF-7, which when disrupted, result in short neuronal dendrites and collapsed sheath channels in the amphid head neurons (Heiman and Shaham, 2009; Low *et al*., 2019). Worms with disrupted TZ gating modules also demonstrate retrograde extension defects, although the abnormalities are mostly restricted to the phasmid pore (tail) (Williams *et al*., 2008, 2011; Williams, Masyukova and Yoder, 2010; Schouteden *et al*., 2015). Two types of retrograde extension defect are described for sensory organs involving glial cell ensheathment. In the first, both the neuronal and sheath cell processes are truncated, and remain associated, as observed for the amphid organ in *dex-1* and *dyf-7* mutants (Heiman and Shaham, 2009). In the second, only the dendrites are short, as reported for several amphid neurons in *grdn-1* mutants, and the phasmid neurons of *mksr-2* and *nphp-4* double mutants (Williams *et al*., 2008; Nechipurenko *et al*., 2016). From our cell junction data (DLG-1::mScarlet), septin mutants may fall into the first class, where connected neuro-glia(sheath) processes detach from where the phasmid pore should form. Indeed, our DLG-1 localisation data indicates that the socket processes may also be short in the septin mutant (**Figures 4C, 5**). As discussed below, the aforementioned genes are thought to serve cell adhesion roles, ensuring proper anchoring of dendritic and glial processes during cell body migration.

### A role for *C. elegans* septins at sensory organ-associated cell junctions

Our prevailing model is that septins regulate phasmid organ morphogenesis and dendritic retrograde extension by facilitating cell adhesive properties at the level of cell junctions (**Figure 5**). In the first instance, we have found that UNC-61 localises at, or very near, DLG-1-marked cell junctions associated with the phasmid organ. These UNC-61 phasmid signals are prominent, and manifest as punctate, short rod and ring-like shapes, precisely in the regions where neuro-glia, glia:glia, and glia:hypodermal/seam adherens junctions are known from fluorescence light and electron microscopy studies (Firestein and Rongo, 2001; Altun and Hall, 2002, 2003). Notably, DLG-1::mScarlet head and tail signals are more prevalent than UNC-61::GFP signals, extending over larger regions; thus, UNC-61 associates with a portion of the DLG-1 signal, indicative of a role within certain types of junction. Importantly, unlike loss-of-function alleles, *unc-61::gfp* knock-in worms lack uncoordinated or gonadal phenotypes, indicating that the GFP tag does not affect *unc-61* function, and that the signals we observe reflect physiological UNC-61 subcellular localisations.

Consistent with our findings, several studies report prominent localisations for mammalian septins (SEPT2/6/9) at epithelial and endothelial cell junctions (Sidhaye *et al*., 2011; Kim and Cooper, 2018, 2021; X. Wang *et al*., 2021; Kim *et al*., 2023; Naydenov *et al*., 2024; Fu *et al*., 2025). Interestingly, as we have observed for UNC-61 and DLG-1, mammalian SEPT2 often localises near, rather than at, sites of core junction components, possibly because septins associate preferentially at membranes with positive curvature (Bridges *et al*., 2016; Kim and Cooper, 2018). Since *unc-61* functions cell non-autonomously to regulate phasmid dendrite extension and cilia, we propose that septins associate with cell-cell junctions in the glia (sheath, socket), a conclusion supported by transcriptomics data showing high levels of *unc-61* expression in these cells (Taylor *et al*., 2021). Notably, several neuro-glia cell junction components (MAGI-1, SAX-7, HMR-1, GRDN-1) were recently reported to cell non-autonomously regulate dendrite extension, specifically in the URX and BAG sensory neurons (head), via a mechanism involving a multivalent adhesion complex of MAGI-1 (glia) and SAX-7 (neuron) (Cebul *et al*., 2024). Indeed, the sensory organ specific nature by which these proteins function correlates with our observations for septins, which regulate dendrite extension in phasmid neurons, but not most examined head neurons, including URX. Thus, dendrite morphogenesis via retrograde extension appears to employ distinct cell junction-associated molecular mechanisms in different sensory organs.

Our finding of reduced DLG-1 levels at the phasmid pore region of *unc-61* mutants further supports a cell junction-associated role for septins in regulating sensory pore morphogenesis. Notably, the affected DLG-1 levels are at the glia(sheath)-glia(socket) and especially the glia(socket)-hypodermis/seam cell junctions, and not the neuro-glia(sheath) junction. We propose that lowered DLG-1 levels indicate loss of adherens junction integrity, which in turn reduces neuro-glia(sheath) process anchoring. In one model, the phasmid sheath:socket and socket-hypodermis/seam cell junctions physically anchor the developing neuro-glia(sheath) processes during retrograde extension to facilitate correct dendrite length and normal cilia length and positioning (**Figure 5**). Alternatively, process adhesion is indirectly facilitated by cell junctions, perhaps via junction-associated signaling that regulates the adhesive properties of the surrounding extracellular matrix. Consistent with this model is that several retrograde extension regulators such as DEX-1 and DYF-7 are secreted (Heiman and Shaham, 2009); also, electron dense DYF-7-dependent extracellular filaments, which may have adhesive properties, extend from the nascent dendritic tips within the developing amphid sensory pore (Oikonomou *et al*., 2011). Whether such filaments exist in the phasmid organ, and whether they might be septin regulated, remains to be shown.

In support of our findings in *C. elegans*, roles for septins in regulating mammalian cell-cell junction organisation have been reported (Kim and Cooper, 2018, 2021; Kim *et al*., 2023; Naydenov *et al*., 2024; Fu *et al*., 2025). Several proposals have been put forward for how septins regulate cell junctions. One idea is that septin filaments closely associated with the plasma membrane stabilise nearby cell junctions, either via direct interactions with junctional proteins or by regulating associated actin assemblies (Zhang *et al*., 1999; Kim and Cooper, 2018; Kim *et al*., 2023). Clearly, more research is required to decipher the general mechanism.

### Conclusions

We have found that *C. elegans* septins regulate a subset of sensory cilia by facilitating neurite retrograde extension during sensory organ morphogenesis. We propose that septins function as a scaffold in glia to stabilise the integrity of adherens cell junctions, which in turn facilitate the necessary cell adhesive properties for sensory organ formation. Given that sense organs across species share conserved structural features and molecular components, septin-dependent junctional regulation represents a broadly relevant mechanism for establishing sensory systems with intact ciliary axonemes. For example, as with *C. elegans* sense organs, sustentacular glia provide structural support for sensory neurons in the vertebrate olfactory epithelium (Gong, 2012; Schwob *et al*., 2017). (Beaudoin, 2016; Naylor *et al*., 2019). Since septin loss affects only a subset of sensory organs in *C. elegans*, more research is required to understand the cell and organ-specific nature of their function, both in worms and higher systems.

## ACKNOWLEDGEMENTS

We thank the University College Dublin Conway Institute imaging facility (Dimitri Scholz, Tiina O’Neill and Niamh Stephens) for assistance with transmission electron microscopy. Research supported by a Taighde Éireann - Research Ireland Government of Ireland Postgraduate scholarship under grant number GOIPG/2021/1572 to EF. For strains, we thank the *C. elegans* genetics center (CGC), funded by an NIH Office of Research Infrastructure Program (P40 OD010440).

---

**Figure S1. Ciliary and dendritic phenotypes for various ciliated head neurons (amphid channel, URX, AWC) in septin disrupted worms. (A)** Representative fluorescence images of the head region of single and double septin gene mutants, dye-filled with DiI. **(B)** Representative fluorescence images of single and double septin gene mutant amphid cilia using a knock-in IFT-74::GFP reporter. Bar; 2 µm. **(C)** Analysis of URX and AWC neuron dendrite morphology and length using *gcy-32*p::*gfp* and *str-2*p*::gfp* reporters. Fluorescence images show the entire neuronal cell structure; the wing-shaped AWC cilia are shown in the smaller panels. Graphs show dendrite length measurements for single and double septin gene mutants. Means denoted by wide horizontal lines. ns; not significant (p>0.05); Mann Whitney test (URX); Kruskal-Wallis test with Dunn’s post-hoc analysis (AWC) vs WT). Scale bars; 5 µm (large panels); 2 µm (small panels).

**Figure S2. UNC-61::GFP subcellular localisation in the head and tail. (A)** Tail images of worms with endogenously tagged UNC-61::GFP (green) showing puncta near the neuronal dendrites stained with DiI (red). **(B)** Airyscan images of the head region of worms with endogenously tagged UNC-61::GFP and the apical cell junction component DLG-1::mScarlet. Scale bars; 10 µm.

**Tables S1-S4**. Strains (S1), primer sequences (S2), CRISPR guide and repair template sequences for *unc-61* and *dlg-1* knockin gfp/mScarlet-tagged alleles (S3) and *unc-61::gfp* transgene construct sequences.

