## Supplementary Material for "*C. elegans* septins regulate a subset of sensory neuronal cilia via cell-non autonomous mechanisms in supporting glia"

### Figure S1

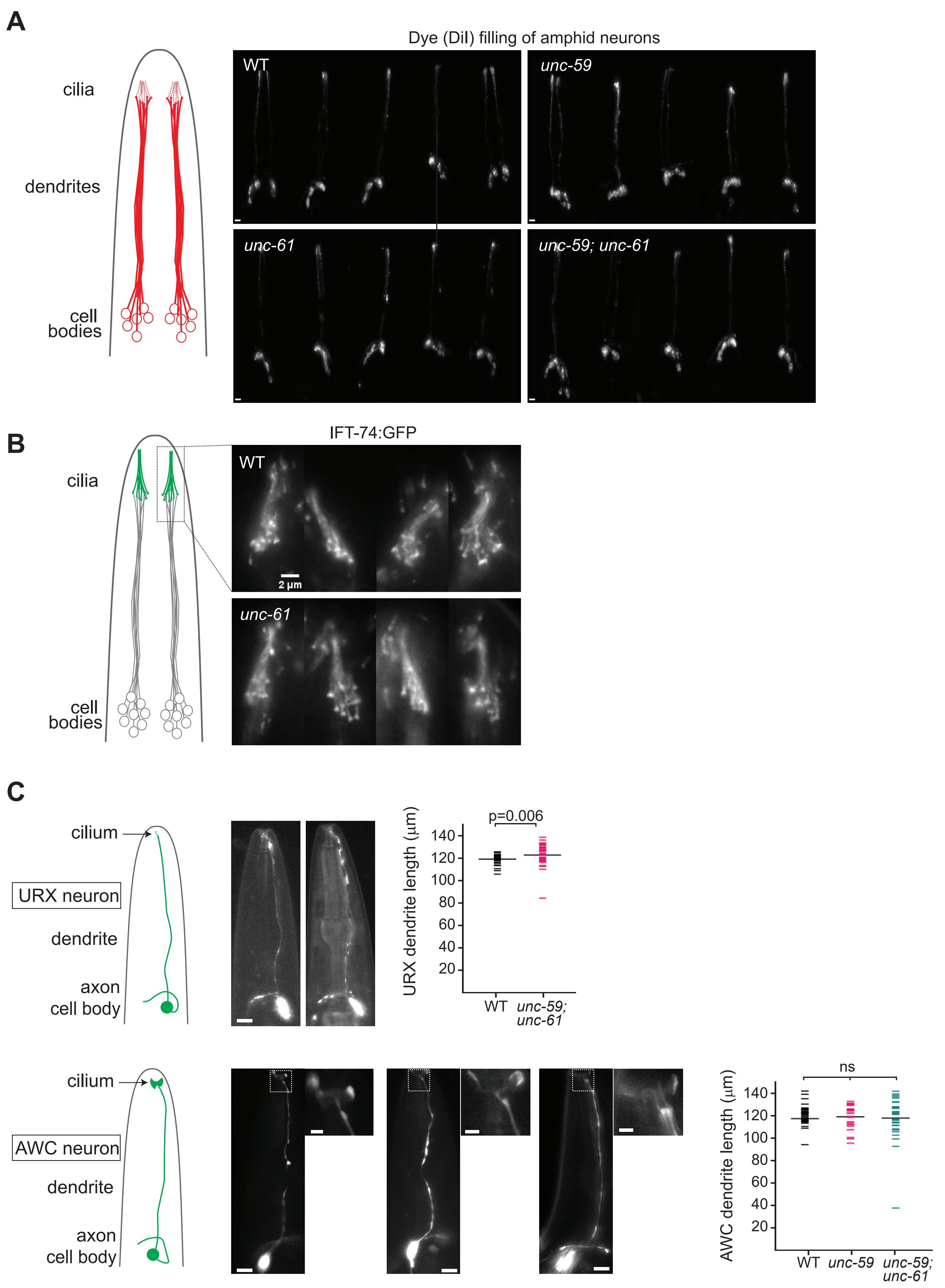
A
Dye (DiI) filling of amphid neurons
cilia
dendrites
cell bodies
WT
unc-59
unc-61
unc-59; unc-61
B
IFT-74:GFP
cilia
cell bodies
WT
2 µm
unc-61
C
cilium
URX neuron
dendrite
axon
cell body
p=0.006
URX dendrite length (µm)
140
120
100
80
60
40
20
WT
unc-59; unc-61
cilium
AWC neuron
dendrite
axon
cell body
ns
AWC dendrite length (µm)
140
120
100
80
60
40
20
WT
unc-59
unc-59; unc-61

### Figure S2

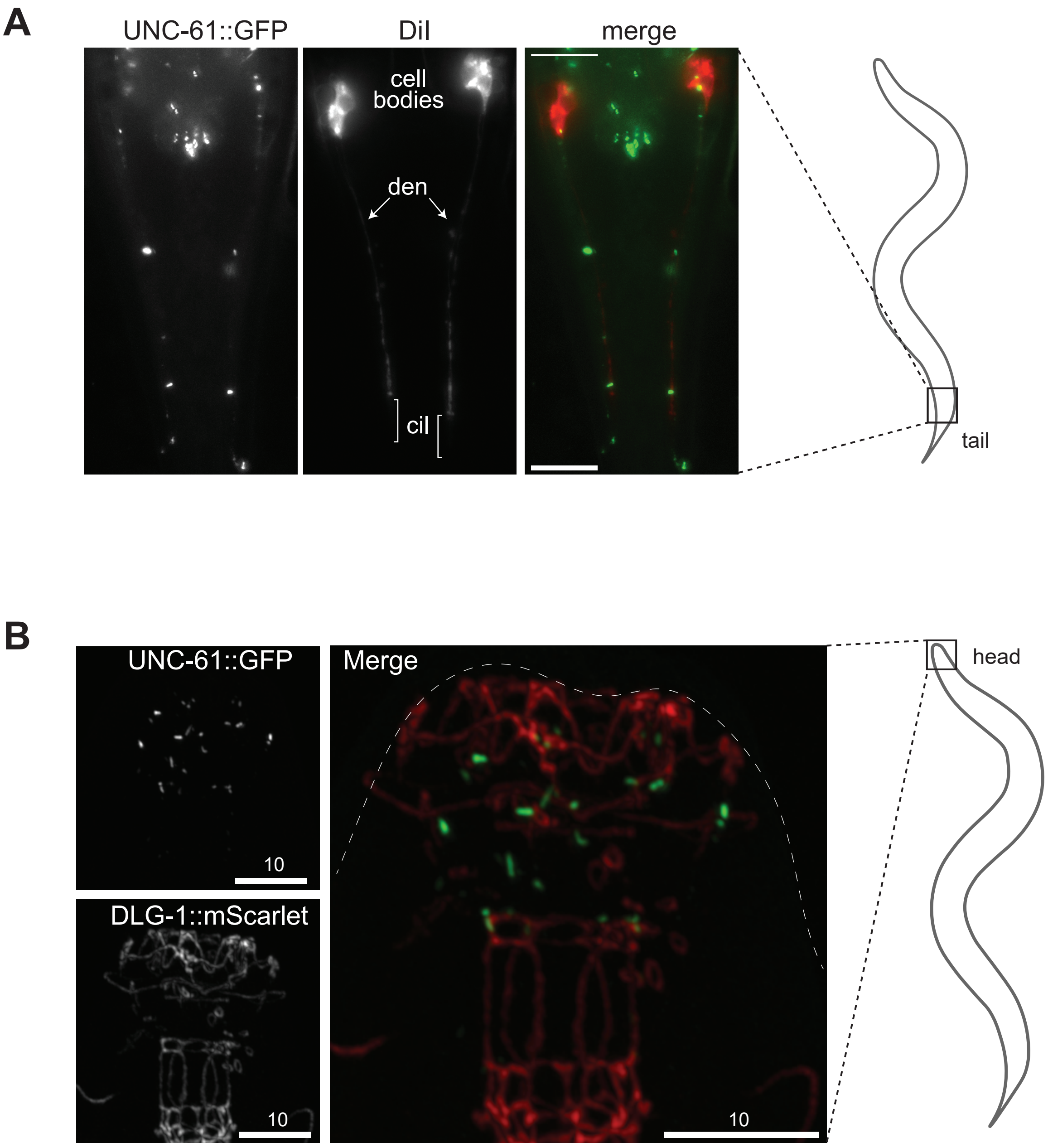
A
UNC-61::GFP
DiI
merge
cell bodies
den
cil
tail
B
UNC-61::GFP
10
DLG-1::mScarlet
10
Merge
10
head

**Table S1. Strains used in this study**

| ID | Genotype | Source |
|---|---|---|
| OEB1083 | *unc-59(tm1939) I; unc-61(e228) V* | This study |
| OEB1156 | *unc-59(tm1939) I; ift-74(cas499[ift-74::gfp]) II* | This study |
| OEB1157 | *unc-59(tm1939) I; ift-74(cas499[ift-74::gfp]) II; unc-61(e228) V* | This study |
| OEB1158 | *ift-74(cas499[ift-74::gfp]) II; unc-61(e228) V* | This study |
| OEB1103 | *unc-59(tm1939) I; oqIs1 [rpi-2::gfp; xbx-1::tdTomato + pRF4] III* | This study |
| OEB1119 | *unc-59(tm1939) I; oqIs1 [rpi-2::gfp; xbx-1::tdTomato + pRF4] III; unc-61(e228) V* | This study |
| OEB1104 | *oqIs1 [rpi-2::gfp; xbx-1::tdTomato + pRF4] III; unc-61(e228) V* | This study |
| OEB1099 | *unc-59(tm1939) I; mNG::nphp-4 V* | This study |
| OEB1100 | *unc-59(tm1939) I; mNG::jbts-14 X* | This study |
| OEB1098 | *unc-59(tm1939) I; unc-61(e228) V; mNG::nphp-4 V* | This study |
| OEB1097 | *unc-59(tm1939 I; unc-61(e228) V; mNG::jbts-14 X* | This study |
| OEB1096 | *unc-59(tm1939) I; mks-6 (gk674) I; unc-61(e228) V* | This study |
| OEB1110 | *unc-59(tm1939) I; nphp-4(tm925) V* | This study |
| OEB1111 | *unc-59(tm1939) I; mks-6 (gk674) I* | This study |
| OEB1213 | *unc-59(tm1939) I; unc-61(e228) V; nphp-4(tm925) V* | This study |
| OEB1181 | *oqEx951 [mir-228*p*::unc-61::gfp + pRF4]* | This study |
| OEB1159 | *oqEx952 [srb-6*p*::unc-61::gfp + pRF4]* | This study |
| OEB1186 | *unc-59(tm1939) I; unc-61(e228) V; kyIs140 [str-2*p*::gfp + lin-15(+)]* | This study |

| | | |
|---|---|---|
| OEB1117 | *unc-59(tm1939) I; kyIs140 [str-2*p*::gfp + lin-15(+)]* | This study |
| OEB1211 | *unc-61(e228) V; oqEx951 [mir-228*p*::gfp::unc::gfp + pRF4]* | This study |
| OEB1212 | *unc-61(e228); oqEx952 [srb-6*p*::unc-61::gfp + pRF4]* | This study |
| OEB1107 | *unc-61(oq173[unc-61::gfp]) V* | This study |
| OEB1232 | *unc-61(oq173[unc-61::gfp]); dlg-1(oq180; [dlg-1::mScarlet]) X* | This study |
| OEB1341 | *ift-74(cas499[ift-74::gfp]) II; unc-61(e228) V;dlg-1(oq180[dlg-1::mScarlet]) X* | This study |
| OEB1342 | *ift-74(cas499[ift-74::gfp]) II; dlg-1(oq180[dlg-1::mScarlet]) X* | This study |
| OEB1344 | *unc-59(tm1939) I; unc-61 (e228) V; iaIs19 [gcy-32*p*::GFP + unc-119(+)]* | This study |
| FX01939 | *unc-59(tm1939) I* | The *C. elegans* Deletion Mutant Consortium et al. (2012) |
| CB288 | *unc-61*(*e228*) V | Brenner (1974) |
| OEB920 | *nphp-4(oq109[mNG::nphp-4) V* | Lange et al. (2021) |
| OEB938 | *jbts-14(oq127[mNG::jbts-14]) X* | Lange et al. (2021) |
| | *nphp-4(tm925) V* | |
| | *mks-6(gk674) I* | |
| GOU2362 | *ift-74(cas499[ift-74::gfp]) II* | Yi P, et al. 2017 |
| OEB1215 | *oqIs1 [pRF4+rpi-2::gfp+xbx-1::tdTomato] III* | formerly *yhEx414*, randomly integrated on Chromosome III |
| CX3695 | *kyIs140 [str-2*p*::gfp + lin-15(+)]* | |
| ZG611 | *iaIs19 [gcy-32*p*::GFP + unc-119(+)]* | |

**Table S2. Primers used in this study**

| Primer ID | Name | Sequence | Purpose |
|---|---|---|---|
| **EF029** | unc59.For+144(tm1939) | CTAATGGTTGTTGgtgaggctc | *unc-59 (tm1939)* genotyping |
| **EF030** | unc59.For-505 | cagacccgcgaagtttgaattag | *unc-59 (tm1939)* genotyping |
| **EF035** | unc59.Rev+687 | gactgtcatggtgccaaatg | *unc-59 (tm1939)* genotyping |
| **EF038** | mks6.For-835 | gcgacaaaggtttgatggacag | *mks-6(gk674)* genotyping |
| **EF039** | mks6.For+1 | gcagaaattcagtttccgttg | *mks-6(gk674)* genotyping |
| **EF040** | mks6.Rev+608 | ctacagaatcttacgcgcatgg | *mks-6(gk674)* genotyping |
| **EF046** | unc61.For+1105ss(e228) | *cgtcgcggaaggtggaataatgtcaaagc* **ttt** | *unc-61 (e228) genotyping (anchor* **allele** foot bridge) see ref. : (Touroutine and Tanis 2020) |
| **EF047** | unc61.For+1105ss(wt) | *cgtcgcggaaggtggaataatgtcaaagc* **ttc** | *unc-61 (e228) genotyping (anchor* **allele** foot bridge) see ref. : (Touroutine and Tanis 2020) |
| **EF070** | unc-61.Rev+1972 | CGAAGAGTTACAAGATCGAGGGC | *unc-61 (e228)* genotyping |
| **EF084** | srb-6p.Rev | TTTTATTTCTTCTGTAGAAATTTC | Fusion PCR to generate *srb-6p::unc-61::gfp* construct |
| **EF085** | srb-6.For-1506 | CTTTGCTGCCCGACCCAATGTGG | Fusion PCR to generate *srb-6p::unc-61::gfp* construct |
| **EF086** | srb-6.For-1469 | cgataggcttctggtattgtg | Fusion PCR to generate *srb-6p::unc-61::gfp* construct |
| **EF091** | unc-61_gfp.For | GAACGTATGAAGTTAATGACTAAG GTCTCCAAGAAGCTCAGAAAGAA GCTTGCATGCCTGCAGGTCGACTC | CRISPR Cas9 of *unc-61::gfp* |
| **EF092** | unc-61_gfp.Rev | GGGATTAATATCAAAATCATAATAC AGTTCGAATAACAGATTCATTTGTA TAGTTCATCCATGCCATGTGTAATC CC | CRISPR Cas9 of *unc-61::gfp* |
| **EF093** | unc-61.Rev+2770 | GGATTAATATCAAAATCATAATAC | CRISPR Cas9 of *unc-61::gfp* |
| **EF094** | unc-61.For+2706.P | [Phos]GAACGTATGAAGTTAATGA C | CRISPR Cas9 of *unc-61::gfp* |
| **EF102** | srb-6p-unc-61.F | GAAATTTCTACAGAAGAAATAAAA ATGAGTTTCGAAACGATTCTC | Fusion PCR to generate *srb-6p::unc-61::gfp* construct |

| | | | |
|---|---|---|---|
| **EF103** | unc-61.Rev+2820 | GGGTGTAGTGTGTATTTGAAATAC | Fusion PCR to generate *srb-6*p::*unc-61::gfp / mir-228*p::*unc-61::gfp* constructs |
| **EF105** | miR-228.F-2196 | TGCAATGCGGGAAGAGACGA | Fusion PCR to generate *mir-228*p::*unc-61::gfp* construct |
| **EF106** | miR-228p.R | ATAAGGAGGAAAATGTCTCGCC | Fusion PCR to generate *mir-228*p::*unc-61::gfp* construct |
| **EF107** | miR-228-unc-61.F | GGCGAGACATTTTCCTCCTTATATGAGTTTCGAAACGATTCTC | Fusion PCR to generate *mir-228*p::*unc-61::gfp* construct |
| **EF114** | mir-228.F-2171 | GATTACTGTATGTGTCGATTTACGG | Fusion PCR to generate *mir-228*p::*unc-61::gfp* construct |
| **EF139** | dlg-1.mScarletKI.Cterm.F | CCATCATCAGCCGTGAATCGCAGACGCCAATTTGGGTGCCACGTCATTTTGGAACCGGAGGTGGCGGATCTG | CRISPR Cas9 of *dlg-1::mScarlet* |
| **EF140** | dlg-1.mScarletKI.Cterm.R | GAAGAAACGATTATTTGTCTAAAAAATATCGAGCTTCATCTACTTGTAGAGCTCGTCCATTCCTCCGGTGGAGTGACG | CRISPR Cas9 of *dlg-1::mScarlet* |
| **EF141** | dlg-1.mScKI.Cterm.R_nested | GAAGAAACGATTATTTGTCTAAAAAAT | CRISPR Cas9 of *dlg-1::mScarlet* |
| **EF143** | dlg-1.mScarlet.P | [Phos]CCATCATCAGCCGTGAATC | CRISPR Cas9 of *dlg-1::mScarlet* |

#### Table S3. CRISPR Cas9 guides and repair templates

| Name | Gene | Location | Sequence |
|---|---|---|---|
| cr-49 | *unc-61* | C-term | TCTTAACTTCTTTGACACTT |
| *unc-61::gfp* repair template | *unc-61* | C-term | GAACGTATGAAGTTAATGACCAAAGTGTCAAAGAAGTTAAGAAAGAAGCTT<br>GCATGCCTGCAGGTCGACTCTAGAGGATCCCCGGGATTGGCCAAAGGACC<br>CAAAGgtatgtttcgaatgatactaacataacatagaacattttcagGAGGACCCTTGAGG<br>GTACCGGTAGAAAAAATGAGTAAAGGAGAAGAACTTTTCACTGGAGTTGT<br>CCCAATTCTTGTTGAATTAGATGGTGATGTTAATGGGCACAAATTTTCTGTCA<br>GTGGAGAGGGTGAAGGTGATGCAACATACGGAAAACTTACCCTTAAATTTA<br>TTTGCACTACTGGAAAACTACCTGTTCCATGGgtaagtttaaacatatatatactaact<br>aaccctgattatttaaattttcagCCAACACTTGTCACTACTTTCTgTTATGGTGTTCA<br>ATGCTTcTCgAGATACCCAGATCATATGAAACgGCATGACTTTTTCAAGAGTG<br>CCATGCCCGAAGGTTATGTACAGGAAAGAACTATATTTTTCAAAGATGACGG<br>GAACTACAAGACACgtaagtttaaacagttcggtactaactaaccatacatatttaaattttca<br>gGTGCTGAAGTCAAGTTTGAAGGTGATACCCTTGTTAATAGAATCGAGTTAA<br>AAGGTATTGATTTTAAAGAAGATGGAAACATTCTTGGACACAAATTGGAATA<br>CAACTATAACTCACACAATGTATACATCATGGCAGACAAACAAAAGAATGGA<br>ATCAAAGTTgtaagtttaaacatgattttactaactaactaatctgatttaaattttcagAACTT<br>CAAAATTAGACACAACATTGAAGATGGAAGCGTTCAACTAGCAGACCATTA<br>TCAACAAAATACTCCAATTGGCGATGGCCCTGTCCTTTTACCAGACAACCAT<br>TACCTGTCCACACAATCTGCCCTTTCGAAAGATCCCAACGAAAAGAGAGAC<br>CACATGGTCCTTCTTGAGTTTGTAACAGCTGCTGGGATTACACATGGCATGG<br>ATGAACTATACAAATGAatctgttattcgaactgtattatgattttgatattaatcc |
| cr-55 | *dlg-1* | C-term | GCCACGTCATTAGatgaaat |
| *dlg-1::mScarlet* repair template | *dlg-1* | C-term | CCATCATCAGCCGTGAATCGCAGACGCCAATTTGGGTGCCACGTCATtttGG<br>AACCGGAGGTGGCGGATCTGGAGGTGGCGGATCCGTCAGCAAGGGAGAG<br>GCAGTTATCAAGGAGTTCATGCGTTTCAAGGTCCACATGGAGGGATCCATG<br>AACGGACACGAGTTCGAGATCGAGGGAGAGGGAGAGGGACGTCCATACG<br>AGGGAACCCAAACCGCCAAGCTCAAGGTCACCAAGGGAGGACCACTCCCA<br>TTCTCCTGGGACATCCTCTCCCCACAATTCATGTACGGATCCCGTGCCTTCAC<br>CAAGCACCCAGCCGACATCCCAGACTACTACAAGCAATCCTTCCCAGAGGG<br>ATTCAAGTGGGAGCGTGTCATGAACTTCGAGGACGGAGGAGCCGTCACCG<br>TCACCCAAGACACCTCCCTCGAGGACGGAACCCTCATCTACAAGGTCAAGC<br>TCCGTGGAACCAACTTCCCACCAGACGGACCAGTCATGCAAAAGAAGACC<br>ATGGGATGGGAGGCCTCCACCGAGCGTCTCTACCCAGAGGACGGAGTCCT<br>CAAGGGAGACATCAAGATGGCCCTCCGTCTCAAGGACGGAGGACGTTACC<br>TCGCCGACTTCAAGACCACCTACAAGGCCAAGAAGCCAGTCCAAATGCCAG<br>GAGCCTACAACGTCGACCGTAAGCTCGACATCACCTCCCACAACGAGGACT<br>ACACCGTCGTCGAGCAATACGAGCGTTCCGAGGGACGTCACTCCACCGGA<br>GGAATGGACGAGCTCTACAAGTAGatgaaattggatatttttagacaaataatcgtt |

#### Table S4. Transgenic construct sequences

| Construct | Sequence |
|---|---|
| *srb-6*p::*unc-61*::*gfp* | cgataggcttctggtattgtgctctttatacttgaaagagcacaacacacctgtgaccagagtcctcatgacaagaatttttttcagtatttgtatatttcgatgagaaaaaacacactttttgtattttt<br>agtttggtcagatctttgccaaaactgctgaacttttgagcaaaactatttgacgcatgactttcattctttgcgattgatgtgctatgcagattttcttgaaactttagcataccgaaattttattaattaa<br>aagcaattcggtcttttcgaaataattttttaaacctcgattaatatatattacttccttttctcccccttttttatgtatgttttatgattgtgatttccaatacgtgtacattcttgattaatttattctctccag<br>caacaagttgcaaattggaattagagagtcagtgttagagaaatacgagtttccctctttttaatgtaaagtttattgtttcaccactcgtgcaggtgttaacatgctttttcgatcgcccttacgaactttt<br>ttatttatatgtcaataaatattttactgatttcataaaattatttttaatcatttatttactattttaaaaaaaaattaaaatgttgcatgattttaataaccaaaattactaattttaaaaagaaacaacact<br>ggttttatgttgtctgatctgagaaattcgttagtgaaatacaagaaaaatagcaaaaataactatgtatgtacaaaataatagggcaccaacaaaaatatttttctgtaatatatttgtcatggttttct<br>ggattttaaaacactcaaaatattatcttgaatgagcaaggcatagcttaatctagagaaaatgatgtgagctgatgtgaacagaaaaaagatacgttccaaaaagtcaccaattttcattgtattatt<br>taccagcattaattttttaaatcaaggtagagtagcgtcagctaggaaattgttaaaacctggataaaattgccagttattataaaaagcatttcaaaaataattttaaattttctaatagtagtcaaa<br>aattggtggttattcagttttgataattcgtaatttaggaaaactaccgtttttttttcaaaattttacaagaagcttttgactagaaattttaaaataactgaaatgttttggagggaatcagtatgaaat<br>ttggtatttttggattattttatatctatttagataaaattggtgccgtttctttgactatttgttgttcaaaagaccagcagaccaaaagctaagcttaaaatttctgcatatatttcagttctttttttacatc<br>gtaaaagaaaaaattctaatctttttacctccattgcgacattttcccagactttatgcatcaaacatacctcctttttttgttttctagaaactactgaaccggatatatgccagtttttattttgtgatcagt<br>cttgaaatttctacagaagaaataaaaATGAGTTTCGAAACGATTCTCTATACTGCATCTGCCCTTCTTTTCATATTTCTATTGGTTCTGCTCACCACCGCTTTTCTCA<br>TCGTCCGACGGTCGAAACAAAGCAGCATAAACCTGCAGACAGTGGTGGTTACCCATGAGAATCCGTATGTCACGATGACCAGTCAATCGTCTAAAATGg<br>tatgaacatctgttgcgcatccattgaaatgacctcgaacatttccaaaaccttgtttaacttctcttttcctcacttacacacatggtttcctccttcttgctaatacatgttttctgaatgcttttctttctct<br>gagtaactcttttatttaaaaaaaaagtctaatttccagGATATCCCAGTGATTCTACCCAAAAGATTCAATATGTCCGACATCGAGCATAAGgtaaattgagaatttttaaacgg<br>tttttcaacaaataattaaagttacagCACTTACCACCTCACCAACCACCTCCACCAGTGCCACATCATCATCAAAACCAGCCAACTCACAATAACACGACTACAAT<br>TTCATCAGCTACGAGCAGCATTAACACCACAACCACGAGCAAGAAGCCAACAATTGCAGCTCCAACGGCTCCTTCACCGATTAAAAGTgtgagtaaattgac<br>tgatttatttggcttaaagctttttagagcgaaccaatcgcattggtaagcccataatcttgccataattatatttttaattatattttctatgttttcagCTCTCAGACCACACTGGCCGTGTTAT<br>GCAACTGAACGGACACGTCGGATTCGATTCTCTTCCTCATCAGCTTGTAAAGAAAGCTGTCGAAGCTGGgtaaccattttcaacattaatttttacttttaaatctaaaa<br>atttcagATTTCAATTCAATCTGATGTGTGTTGGAGAAACGGGAACAGGAAAAACAACACTTATAGAGTCTTTGTTCAACATGAAGCTCGATTTCGAGCCA<br>TGCAATCATGAATTGAAAACTGTTGAGCTGAGAACTTGCACGAAAGgtgagcaacgaaaagtctactgaaacagaaattttgaaattaaatttcagACGTCGCGGAAGG<br>TGGAATACGAGTGAAGCTTCGACTCGTTGAAACTGCCGGATTTGGAGATCAGCTGGATAAAGATAAAAGgtttttatttcagagattttgcgatttttgcgtatccggtc<br>tcgaaacgacaagtttcattgtttttcaaagtttttcaattaaaaaaaaagttttatttatttattaaaagctcattcaacaatacacttgaactataacaaattgtgagaaaactacgaaattttataa<br>aaattccgcagcaacgaatatttggaattacagtaatcattgaaagcgcacacttctcgcatttaacaaaaaattgtcgtgttgagaacgggaaccgtatttttgcagcaaaaatcgcaaaatgatgc<br>gtccgggtgataaaaaagttatgatctgaaaggaaaaataataattttcagTGCCAAAGTAATTGTCGATTATCTCGAATCGCAGTTTGAAACATACCTCCAAGAAGAGTT<br>GAAGCCACGTCGAATGCTTCAGTATTTCAATGATTCGAGAATCCACGCATGTCTCTACTTCATATCACCTACTGGGCACGGgtaggtggaatatactctgaaatta<br>agtttcagagggttttcagagggttaagctacaaagcttctatttaaaaccgcggtgtgcggcaacaattttgtttttaaaataacatttatttcggctaaaccaattgaaaaatgcacaacatttccaa<br>ctcagagttggaaatgccatagaaatttcctgcaattttcaactttcaggtgatttttgagcatttttcaatcattttaactgaaatcaaaactgttttgtagaattgccgcgctttcctgatttcaaactcta<br>aattgtaactaattttatctaatctttcagACTCAAAGCCCTCGATCTTGTAACTCTTCGCGAATTGGCTAAGCGTGTTAATGTGATCCCAGTGATAGCGAAATCAGA<br>CACAACTTGCAAGGATGAGCTTCTCAGATTCAAAGCGAAAATATTGAGCGAGCTGAAATCTCAGAAAATCGATATTTACACGTTCCCAACTGACGATGA<br>AACTGTCTCAACGACGAACAAGGAAATGAATAAATCGGTTCCGTTCGCCGTTGTTGGAAGTATTGATTTTGTGAAGAAAGAGAATGGACAAATGGTTC<br>GTGCTCGTCAATATCCATGGGGAATAGTCGAAGTGGAGAATGAATCACATTGTGATTTTGTTAAACTCCGTGAAGCACTTCTACGTACAAATGTTGATGA<br>GATGAGACAACGAACTCACGAGTCACTCTATGAAAATTACCGTCGTGACAGACTTCGTCAGATGAAGATTGGAGATGGAGAGACTGGACCAAAGATTA<br>TTGAAAAACTCGCACAGgtattatttcagagaaaatattgtttcgtttcactgattctaagcgtcgattgcagAAACATCGCGAGCATCAAGACGAGTTCAGCCGTCGTGAG<br>CTTACTCTTCGCGAAGAATTTCAGAAGAAGCTTGATGTAACAGAAGGTGACATGAGAAAAGTTGAAGAAGGATTGGCTGCACGCGAGCGAGAGGTTC<br>ATGAGAATTATAATCGAGAGGCGTCGAAACTTGATATGGAGATTCGTCAATTGACTGAAGAACGTATGAAGTTAATGACCAAAGTGTCAAAGAAGTTAA<br>GAAAGAAGCTTGCATGCCTGCAGGTCGACTCTAGAGGATCCCCGGGATTGGCCAAAGGACCCAAAGgtatgtttcgaatgatactaacataacatagaacattttca<br>gGAGGACCCTTGAGGGTACCGGTAGAAAAAATGAGTAAAGGAGAAGAACTTTTCACTGGAGTTGTCCCAATTCTTGTTGAATTAGATGGTGATGTTAAT<br>GGGCACAAATTTTCTGTCAGTGGAGAGGGTGAAGGTGATGCAACATACGGAAAACTTACCCTTAAATTTATTTGCACTACTGGAAAACTACCTGTTCCA<br>TGGgtaagtttaaacatatatatactaactaaccctgattatttaaattttcagCCAACACTTGTCACTACTTTCTgTTATGGTGTTCAATGCTTcTCgAGATACCCAGATCATA<br>TGAAACgGCATGACTTTTTCAAGAGTGCCATGCCCGAAGGTTATGTACAGGAAAGAACTATATTTTTCAAAGATGACGGGAACTACAAGACACgtaagttt<br>aaacagttcggtactaactaaccatacatatttaaattttcagGTGCTGAAGTCAAGTTTGAAGGTGATACCCTTGTTAATAGAATCGAGTTAAAAGGTATTGATTTTAA<br>AGAAGATGGAAACATTCTTGGACACAAATTGGAATACAACTATAACTCACACAATGTATACATCATGGCAGACAAACAAAAGAATGGAATCAAAGTTgta<br>agtttaaacatgattttactaactaactaatctgatttaaattttcagAACTTCAAAATTAGACACAACATTGAAGATGGAAGCGTTCAACTAGCAGACCATTATCAACAA<br>AATACTCCAATTGGCGATGGCCCTGTCCTTTTACCAGACAACCATTACCTGTCCACACAATCTGCCCTTTCGAAAGATCCCAACGAAAAGAGAGACCACA<br>TGGTCCTTCTTGAGTTTGTAACAGCTGCTGGGATTACACATGGCATGGATGAACTATACAAATGAatctgttattcgaactgtattatgattttgatattaatcccattttgtt<br>ttatttccgtttttttatgtatttcaaatacacactacaccc |

| | |
|---|---|
| *mir-228*p::*unc-61*::*gfp* | gattactgtatgtgtcgatttacggcattaattttgagtttgcatgaatccatatatatatatatatatatctcttttgtaaaagtttctgtgttgagtttgcattgtccaatgtttacttaaattagttaaataa<br>tttagttaaataatcctgatgaaactgtaattttactaatcggtgaccctggctgaaaaatgtttattgtgacgtcatactcttgcaaaaacggttgcattgctccgatatggacaacttcccagagcagt<br>gcggtttttgcaagatagcctaaaaagagattcatattcgatttttccctttagacctttccgaagctggccgacaattgtcttgaaaggggtttccaagggataggtggccgagtttttgtattccgcga<br>acacatattaccgttttgggaacttgatagccggcgtcaatttaagaatacgcggaataaaacataaatcagatattatcctacgttaattcaaaatatttttgaaatttgtttaatgtttcagttcaaat<br>ttatgtatctttcgcaaacgtttctgtagttcctttttgctttcgtggaaggattcattttgattgtaaaaaacaaacaaagtttcatcggttagaactccaattaaaatatttttaccatttcccacattata<br>ggaacataagtattcttattttgaagttactttcttggagagtaattaacgggaaaccttatgcgtaaaattttcaaaaattagcatgtttcaattgataagactcaccggataacaaaactatcagtag<br>aattaattaatatgaatgtttttaaaatatgcatgaaacagttttttctcaattcaaaaattattctgacttaatcaagtcataatatttagcataaatattaagtacacctggaacagaagtaacatta<br>atcgtagtttttttgacagaaatgtaccatcgaaaatgattacataaacatttattgtaacatttgagttagttaactggacctaatgaaactgaacattatggcagtgagtctaatgaattttggtctca<br>aactaccaaaagttttcttatattcaggaataccacaaaaaaaacattaaaaactccacatccacgagagttgtattctcaggtagtttcaaattaaactttttctttgatagttttggtcttaaattcatc<br>ggactcactgccattatagacattatttattcggtttttttaaagaatttattaaagaactacaatatatgtttcttttattgttggtggaagtttgcgaatttttagactagcagatcctcatctggtaatttt<br>taaaatttacttccacaaactaatttattttgctgttctcacgtatatagaaaataatttaaaaattaaatttgaattcccgcgtaaaaccatttgcgtgcgaaattcaaattcagccactttcgtcgatttt<br>aatggcgaaattttcattggtgacggatgttcccgtttttaaattgttttcatagttgataagctaaggacttaataataagtattttcccttttgaggccttcttaattatgttctgaaacagtagaaata<br>agagattccaccataaatcgtgttcaattcggataatccttgtgtgttcattttcatgacccttcatagtgagggaataattggtctctctctctctctctctctctctctctctctctctctctctttttct<br>ctaaaactctctcaatcccttacttttcaacgacctcgacagtatttcaaatgtacacagaacaatagaaatcgcctcgtttctgtttctctctctctctctctctctctctctctctctcttacttcttccaa<br>ttgaatcgacaattttcgaaatcgaaataggccacagtgaggaaggccgacgaattgagggaaatacacagaaaccccccgccccataatctgtagagaaaaaagagagaaatgggcggagctt<br>gcttactcttttggcgcctccttctctttctctaagtctcactttccatctgttctttcttctccacatacagtcattcgttggcctgtgctgcctcgtcgactactactactgccttcaaaaattcttttattcgt<br>cgcctcccaaaaactgtttcgcaacttttctattttttttgttgctttggcgagacattttcctccttatATGAGTTTCGAAACGATTCTCTATACTGCATCTGCCCTTCTTTTCATATTT<br>CTATTGGTTCTGCTCACCACCGCTTTTCTCATCGTCCGACGGTCGAAACAAAGCAGCATAAACCTGCAGACAGTGGTGGTTACCCATGAGAATCCGTATG<br>TCACGATGACCAGTCAATCGTCTAAAATGgtatgaacatctgttgcgcatccattgaaatgacctcgaacatttccaaaaccttgtttaacttctcttttcctcacttacacacatggtttcct<br>ccttcttgctaatacatgttttctgaatgcttttctttctctgagtaactcttttatttaaaaaaaagtctaatttccagGATATCCCAGTGATTCTACCCAAAAGATTCAATATGTCCGAC<br>ATCGAGCATAAGgtaaattgagaatttttaaacggtttttcaacaaataattaaagttacagCACTTACCACCTCACCAACCACCTCCACCAGTGCCACATCATCATCAAAA<br>CCAGCCAACTCACAATAACACGACTACAATTTCATCAGCTACGAGCAGCATTAACACCACAACCACGAGCAAGAAGCCAACAATTGCAGCTCCAACGG<br>CTCCTTCACCGATTAAAAGTgtgagtaaattgactgatttatttggcttaaagctttttagagcgaaccaatcgcattggtaagcccataatcttgccataattatatttttaattatattttctatg<br>ttttcagCTCTCAGACCACACTGGCCGTGTTATGCAACTGAACGGACACGTCGGATTCGATTCTCTTCCTCATCAGCTTGTAAAGAAAGCTGTCGAAGCTG<br>GgtaaccattttcaacattaatttttacttttaaatctaaaaatttcagATTTCAATTCAATCTGATGTGTGTTGGAGAAACGGGAACAGGAAAAACAACACTTATAGAGT<br>CTTTGTTCAACATGAAGCTCGATTTCGAGCCATGCAATCATGAATTGAAAACTGTTGAGCTGAGAACTTGCACGAAAGgtgagcaacgaaaagtctactgaaac<br>agaaattttgaaattaaatttcagACGTCGCGGAAGGTGGAATACGAGTGAAGCTTCGACTCGTTGAAACTGCCGGATTTGGAGATCAGCTGGATAAAGATAA<br>AAGgtttttatttcagagattttgcgatttttgcgtatccggtctcgaaacgacaagtttcattgtttttttcaaagtttttcaattaaaaaaaaagttttatttatttattaaaagctcattcaacaatacact<br>tgaactataacaaattgtgagaaaactacgaaattttataaaaattccgcagcaacgaatatttggaattacagtaatcattgaaagcgcacacttctcgcatttaacaaaaaattgtcgtgttgaga<br>acgggaaccgtatttttgcagcaaaaatcgcaaaatgatgcgtccgggtgataaaaaagttatgatctgaaaggaaaaataataattttcagTGCCAAAGTAATTGTCGATTATCTCGAA<br>TCGCAGTTTGAAACATACCTCCAAGAAGAGTTGAAGCCACGTCGAATGCTTCAGTATTTCAATGATTCGAGAATCCACGCATGTCTCTACTTCATATCACC<br>TACTGGGCACGGgtaggtggaatatactctgaaattaagtttcagagggttttcagagggttaagctacaaagcttctatttaaaaccgcggtgtgcggcaacaattttgtttttaaaataacatt<br>tatttcggctaaaccaattgaaaaatgcacaacatttccaactcagagttggaaatgccatagaaatttcctgcaattttcaactttcaggtgatttttgagcatttttcaatcattttaactgaaatcaa<br>aactgttttgtagaattgccgcgctttcctgatttcaaactctaaattgtaactaattttatctaatctttcagACTCAAAGCCCTCGATCTTGTAACTCTTCGCGAATTGGCTAAGCG<br>TGTTAATGTGATCCCAGTGATAGCGAAATCAGACACAACTTGCAAGGATGAGCTTCTCAGATTCAAAGCGAAAATATTGAGCGAGCTGAAATCTCAGAA<br>AATCGATATTTACACGTTCCCAACTGACGATGAAACTGTCTCAACGACGAACAAGGAAATGAATAAATCGGTTCCGTTCGCCGTTGTTGGAAGTATTGAT<br>TTTGTGAAGAAAGAGAATGGACAAATGGTTCGTGCTCGTCAATATCCATGGGGAATAGTCGAAGTGGAGAATGAATCACATTGTGATTTTGTTAAACTC<br>CGTGAAGCACTTCTACGTACAAATGTTGATGAGATGAGACAACGAACTCACGAGTCACTCTATGAAAATTACCGTCGTGACAGACTTCGTCAGATGAAG<br>ATTGGAGATGGAGAGACTGGACCAAAGATTATTGAAAAACTCGCACAGgtattatttcagagaaaatattgtttcgtttcactgattctaagcgtcgattgcagAAACATCGC<br>GAGCATCAAGACGAGTTCAGCCGTCGTGAGCTTACTCTTCGCGAAGAATTTCAGAAGAAGCTTGATGTAACAGAAGGTGACATGAGAAAAGTTGAAG<br>AAGGATTGGCTGCACGCGAGCGAGAGGTTCATGAGAATTATAATCGAGAGGCGTCGAAACTTGATATGGAGATTCGTCAATTGACTGAAGAACGTATG<br>AAGTTAATGACCAAAGTGTCAAAGAAGTTAAGAAAGAAGCTTGCATGCCTGCAGGTCGACTCTAGAGGATCCCCGGGATTGGCCAAAGGACCCAAAG<br>gtatgtttcgaatgatactaacataacatagaacattttcagGAGGACCCTTGAGGGTACCGGTAGAAAAAATGAGTAAAGGAGAAGAACTTTTCACTGGAGTTGTC<br>CCAATTCTTGTTGAATTAGATGGTGATGTTAATGGGCACAAATTTTCTGTCAGTGGAGAGGGTGAAGGTGATGCAACATACGGAAAACTTACCCTTAAA<br>TTTATTTGCACTACTGGAAAACTACCTGTTCCATGGgtaagtttaaacatatatatactaactaaccctgattatttaaattttcagCCAACACTTGTCACTACTTTCTgTTATG<br>GTGTTCAATGCTTcTCgAGATACCCAGATCATATGAAACgGCATGACTTTTTCAAGAGTGCCATGCCCGAAGGTTATGTACAGGAAAGAACTATATTTTTC<br>AAAGATGACGGGAACTACAAGACACgtaagtttaaacagttcggtactaactaaccatacatatttaaattttcagGTGCTGAAGTCAAGTTTGAAGGTGATACCCTTGTT<br>AATAGAATCGAGTTAAAAGGTATTGATTTTAAAGAAGATGGAAACATTCTTGGACACAAATTGGAATACAACTATAACTCACACAATGTATACATCATGGC<br>AGACAAACAAAAGAATGGAATCAAAGTTgtaagtttaaacatgattttactaactaactaatctgatttaaattttcagAACTTCAAAATTAGACACAACATTGAAGATGG<br>AAGCGTTCAACTAGCAGACCATTATCAACAAAATACTCCAATTGGCGATGGCCCTGTCCTTTTACCAGACAACCATTACCTGTCCACACAATCTGCCCTTT<br>CGAAAGATCCCAACGAAAAGAGAGACCACATGGTCCTTCTTGAGTTTGTAACAGCTGCTGGGATTACACATGGCATGGATGAACTATACAAATGAatctg<br>ttattcgaactgtattatgattttgatattaatcccattttgttttatttccgtttttatgtatttcaaatacacactacaccc |
